# Adaptation to chronic ER stress enforces pancreatic β-cell plasticity

**DOI:** 10.1101/2021.05.24.445193

**Authors:** Chien-Wen Chen, Bo-Jhih Guan, Mohammed R. Alzahrani, Zhaofeng Gao, Long Gao, Syrena Bracey, Leena Haataja, Ashleigh E. Schaffer, Hugo Lee, Thomas Laframboise, Ilya Bederman, Peter Arvan, Clayton E. Mathews, Ivan C. Gerling, Klaus H. Kaestner, Boaz Tirosh, Feyza Engin, Maria Hatzoglou

**Author notes:** Co-corresponding (C-W.C.), (F.E.), (M.H.).

## Abstract

Pancreatic β-cells undergo high levels of endoplasmic reticulum (ER) stress due to their role in insulin secretion. Hence, they require sustainable and efficient adaptive stress responses to cope with the stress. Whether duration and episodes of chronic ER stress directly compromises β-cell identity is largely unknown. We show that under reversible, chronic ER stress, β-cells undergo a distinct transcriptional and translational reprogramming. During reprogramming, expression of master regulators of β-cell function and identity and proinsulin processing is impaired. Upon recovery from stress, β-cells regain their identity, highlighting a high-degree of adaptive β-cell plasticity. Remarkably, when stress episodes exceed a certain threshold, β-cell identity is gradually lost. Single cell RNA-seq analysis of islets from type 1 diabetes (T1D) patients, identifies the severe deregulation of the chronic stress-adaptation program, and reveals novel biomarkers for progression of T1D. Our results suggest β-cell adaptive exhaustion (βEAR) is a significant component of the pathogenesis of T1D.

## INTRODUCTION

Pancreatic β-cells have a highly active secretory pathway, and may reach a production rate as high as 1 million molecules of pre-proinsulin per minute under stimulated conditions (Scheuner and Kaufman, 2008), which is a physiological ER stress. Therefore, tolerance to ER stress is an integral component of the normal physiology of β-cells, as demonstrated by their well-developed and effective adaptive stress response mechanisms (Eizirik and Cnop, 2010). In addition to physiological stress, several pathological conditions can elicit uncompensated, severe ER stress in β-cells, including reactive oxygen species, toxins, viral infections and inflammation (Fonseca et al., 2011). All of these conditions trigger the unfolded protein response (UPR), which is mediated by the coordinated actions of ER membrane-localized inositol-requiring protein 1 alpha (IRE1α), activating transcription factor 6 (ATF6) and the eIF2α PKR-like endoplasmic reticulum kinase (PERK) (Ron and Walter, 2007). Under acute ER stress, the adaptive UPR aims to establish a return to cellular homeostasis, while chronic and unresolvable stress can result in a maladaptive UPR that contributes to various pathologies, including diabetes (Krokowski et al., 2013; Walter and Ron, 2011). There is a time frame of physiological stress during which β-cells induce adaptive compensatory mechanisms (Stamateris et al., 2013).

While IRE1α activity has been implicated in the transition from adaptive to maladaptive UPR (Han et al., 2009; Morita et al., 2017), the impact of recovery from chronic severe stress on β-cell identity has not been well-defined. Several lines of evidence implicate ER stress and aberrant UPR in the pathogenesis of both type 1 and type 2 diabetes (T1D and T2D, respectively) (Arunagiri et al., 2018; Arunagiri et al., 2019; Back and Kaufman, 2012; Eizirik et al., 2020; Engin, 2016; Engin et al., 2014; Lee et al., 2020; Morita et al., 2017; Ozcan et al., 2004; Sims et al., 2020). Indeed, mutations in genes that influence the ER stress responses frequently result in loss of β-cell function and/or death in both experimental models and in humans, including the genes encoding the major UPR elements PERK, eIF2α, CHOP, XBP1, IRE1α, and WSF1 (Fonseca et al., 2011; Harding et al., 2001; Zhang and Kaufman, 2006). In this study, we investigated the molecular underpinnings of reversible chronic and episodic ER stress and their effects on β-cell adaptation and plasticity.

The use of chemical stress is critical to observe temporal changes during progression from acute to chronic ER stress in cells (Guan et al., 2017) and *in vivo* (Abdullahi et al., 2017; Gomez and Rutkowski, 2016; Schneider et al., 2020; Zhang et al., 2014). Genetic chronic ER stress models (Bommiasamy and Popko, 2011; Yoshioka et al., 1997; You et al., 2021) do not provide the synchronized control over these transitions; however, numerous studies show that the biochemical principles of the physiological ER stress response are shared with chemically-induced ER stress *in vitro* and *in vivo* (Gomez and Rutkowski, 2016; Schneider et al., 2020; Zhang et al., 2014). Therefore, we selected the reversible inhibitor of the sarcoendoplasmic reticulum pump Ca^++^ ATPase (SERCA) pump, cyclopiazonic acid (CPA) to model resolvable chronic ER stress in the mouse insulinoma cell line, MIN6. This inhibitor induces bona fide ER stress that can be carefully tuned in duration by washing it out of the cells and thereafter monitoring recovery from stress (Guan et al., 2017). Using CPA, we showed that β-cells undergo a homeostatic state change associated with an adaptive reprogramming of their transcriptome and translatome that compromises β-cell identity. A comparative analysis of adaptive reprogramming between β-cells (MIN6) and mouse embryonic fibroblasts (MEFs), revealed an elaborate, β-cell specific network of genes that exhibit stress-induced changes in their expression. Adaptation to chronic ER stress in β-cells included 334 genes, with a predominant downregulation of genes involved in Maturity Onset Diabetes of the Young (MODY), proinsulin expression, synthesis, processing and secretion. This was in contrast to MEFs, which did not show differential gene expression for these genes expressed in both cell types during chronic ER stress (Guan et al., 2017). In addition, β-cells induced more than half of the genes known to participate in ER protein processing, 133 under chronic adaptation conditions. We call this subset of genes the β-cell-specific ER adaptosome. Upon relief from stress, β-cells were gradually able to regain expression of mature identity markers. Remarkably, upon iterative cycles of ER stress, β-cells lost plasticity and expression of their identity genes progressively declined. Laser capture microdisected islets (Holm et al., 2018; Trembath et al., 2020) and single-cell RNA sequencing (scRNA-seq) data from patients with T1D (PancDB; https://hpap.pmacs.upenn.edu) found gene expression changes in β-cells of patients with T1D, in agreement with chronic ER stress adaptive exhaustion and deregulation of the β-cell ER adaptosome that correlated with disease progression. We propose that cycles of ER stress exhaust the β-cell’s adaptive plasticity, and promotes an irreversible loss of function that likely contributes to T1D pathogenesis. We call this novel mechanism the ‘<u>β</u>-cell <u>e</u>xhaustive <u>a</u>daptive response (βEAR)’.

## Results

### Adaptation to chronic ER stress in MIN6 cells sustains proinsulin and insulin protein levels

MIN6 cells are a mouse insulinoma β cell line, isolated from C57BL/6, that displays characteristics of β-cells, including insulin biosynthesis and secretion in response to glucose (Ishihara et al., 1993; Miyazaki et al., 1990). We tested the effect of severe ER stress on the ability of these cells to survive as well as on proinsulin synthesis. MIN6 cells were treated with CPA (Moncoq et al., 2007), which disrupts Ca^++^ homeostasis in the ER, generating an unfavorable protein folding environment that induces the UPR (Lytton et al., 1991).

As we observed previously in MEFs (Guan et al., 2017), CPA treatment induced a time-dependent ER stress response in MIN6 cells. Phosphorylation of PERK and its downstream target, the translation initiation factor eIF2α, was increased one hour following CPA exposure, and was maintained for the duration of treatment (24h), indicative of a chronic ER stress condition. As expected, the expression of hallmark proteins of the PERK branch of the UPR activation, ATF4 and GADD34, showed a gradual increase from 1h to 24h of treatment (**Fig. 1A**), consistent with the negative feedback mechanism of protein synthesis recovery downstream of ATF4 (Novoa et al., 2003).

**Figure 1.**
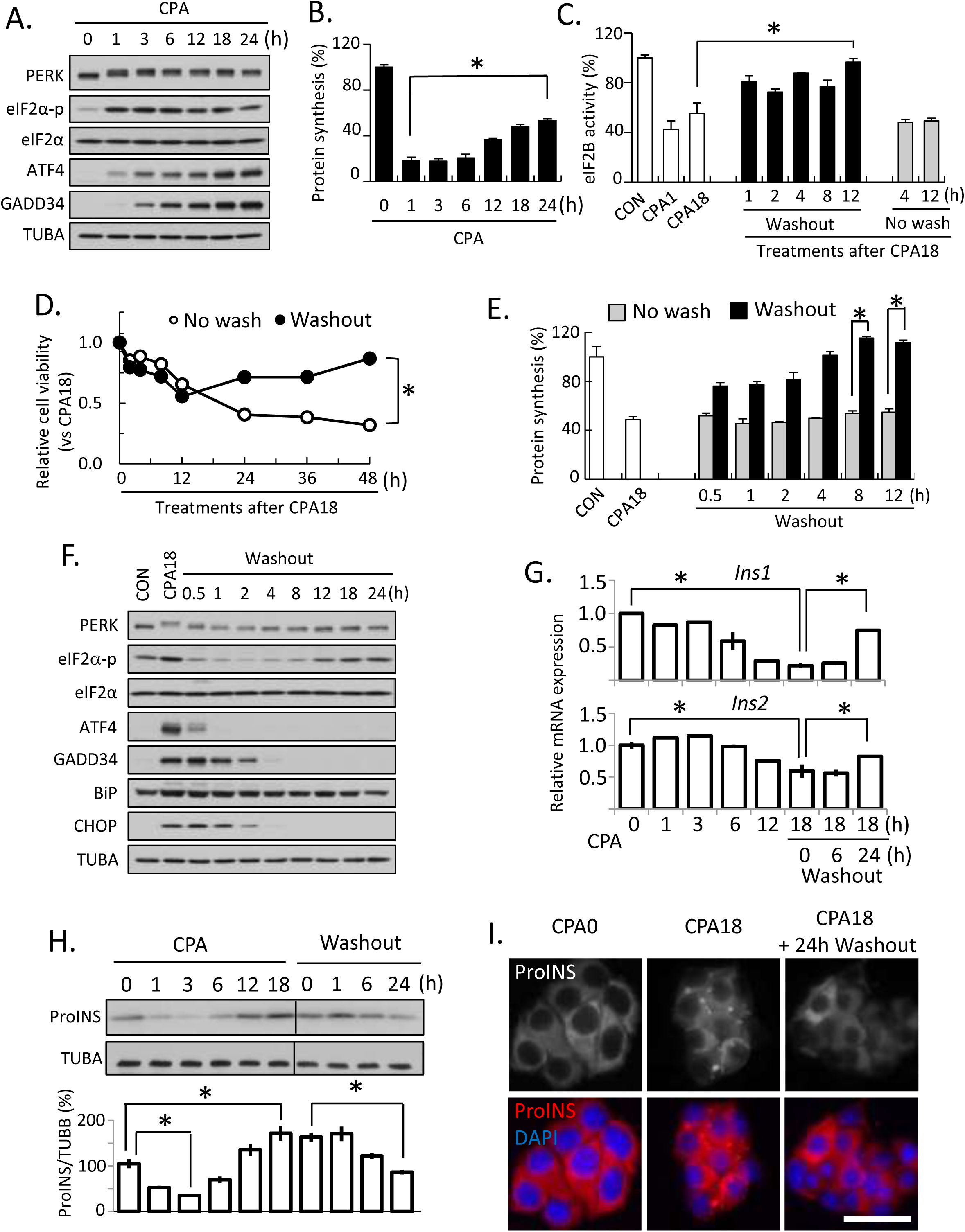
MIN6 cells can simultaneously survive chronic ER stress and alter proinsulin proteostasis. Western blot analysis (A), protein synthesis (B), and eIF2B GEF activity (C), measured in MIN6 cells treated with CPA as indicated. Cell viability (D), protein synthesis (E), and Western blot analysis (F) measured in MIN6 cells for the indicated treatments. qRT-PCR analysis for *Ins1 and Ins2* mRNA levels normalized to *GAPDH* (G), and Western blot analysis for proinsulin (H), measured in MIN6 cells for the indicated treatments. Fluorescence immunocytochemistry of proinsulin at 0 and 18h of CPA treatment and 24h of washout following CPA treatment. Scale bar represents 50 μm. Error bars represent S.E.M. * p<0.05 two-tailed paired Student’s t-test. Representative western blotting images were shown.

MEFs show a dramatic decrease of protein synthesis during the early response to ER stress, which partially recovers during the chronic phase (Guan et al., 2017). The inhibition of protein synthesis is due to the repressive function of phosphorylated eIF2α (eIF2α-p) on the activity of the guanine nucleotide exchange factor eIF2B, an essential translation initiation factor for delivery of the initiator met-tRNA_i_ to the AUG initiation codons of mRNAs (Ron, 2002; Sood et al., 2000). In MEFs, the recovery of protein synthesis was independent of restored eIF2B activity, which remains inhibited during chronic ER stress (Guan et al., 2017). Similarly to situation in MEFs, MIN6 cells showed inhibition of protein synthesis followed by partial recovery (**Fig. 1B**), without recovery of eIF2B activity (**Fig. 1C**). These data indicate that MIN6 cells behave similar to MEFs in adapting to chronic ER stress.

In order to determine the prosurvival adaptive features of the chronic ER stress response in MIN6 cells, we tested the viability of the cells which were maintained in CPA or when they were incubated in normal media following 18h of CPA-exposure. The resumption of growth following recovery from 18h of ER stress confirmed the adaptive chronic ER stress response at that timepoint (**Fig. 1D**). The ability of MIN6 cells to recover from chronic ER stress was also accompanied by biochemical indicators of protein synthesis recovery (**Fig. 1E**), and restoration of eIF2B activity (**Fig. 1C**). Similarly, relief from ER stress inactivated the stress-response signaling and stress-induced gene expression (**Fig. 1F**), suggesting a return to homeostasis (Fumagalli et al., 2016; Guan et al., 2017). Taken together, we conclude that MIN6 cells have the same core mechanism of regulating global protein synthesis rates during adaptation to chronic ER stress as observed in MEFs (Guan et al., 2017).

In MEFs, we have shown that adaptive translational reprogramming involves an eIF4E-dependent to an eIF4E-independent, eIF3-dependent switch of mRNA translation (Guan et al., 2017). Because the most efficient mechanism of mRNA translation initiation utilizes eIF4F, of which eIF4E is the core component (Pestova et al., 2001), we proposed that this switch allows inefficient translation of all mRNAs and more specifically ER-associated mRNAs (Guan et al., 2017). However, in MEFs, we also found that a subset of mRNAs, including transcripts encoding for stress-induced and ER synthesized-proteins, are translated during adaptive chronic ER stress as part of the reprogrammed translation initiation mechanism (Guan et al., 2017). Consistent with MEFs (Guan et al., 2017), accumulation of stress-induced proteins in MIN6 cells exposed to CPA, was similar in control and eIF4E-depleted cells (**Sup. Fig. 1A**), further supporting the adaptive, eIF4E-independent reprogramming of β-cells during chronic ER stress.

Because MIN6 cells are highly specialized insulin secreting cells, we next determined, the regulation of proinsulin gene (*Ins1* and *Ins2*) expression during the course of transitioning from the acute to chronic ER stress caused by CPA treatment. Levels of the *Ins1* and *Ins2* mRNAs decreased in MIN6 cells during chronic ER stress, suggesting an adaptive response of the cells to limit ER mRNA load for the abundant ER synthesized proteins. This decrease was gradually reversed during recovery from stress (**Fig. 1G**), further supporting the regulation of proinsulin gene expression as part of the adaptive response to chronic ER stress. We next examined the levels of both mature insulin and proinsulin under chronic adaptive ER stress and recovery. Neither the insulin monomer nor native insulin hexamers were altered in abundance during ER stress and recovery (**Sup. Fig. 1B**). In contrast to the significantly decreased levels of *Ins2* mRNA, during adaptive chronic ER stress, proinsulin protein levels decreased dramatically early in the stress response, presumably due to eIF2α-p-dependent inhibition of the ER-associated proinsulin mRNA translation (Reid et al., 2014). Subsequently, proinsulin protein levels returned to normal levels at ≥12h despite decreased transcript levels during the chronic adaptive stress period (**Fig. 1H**). Upon recovery from chronic stress, proinsulin protein levels remained similar to that of the untreated control (**Fig. 1H**). These data suggest that proinsulin mRNA participates in the adaptive reprogramming of mRNA translation during chronic ER stress. Finally, in order to determine if proinsulin processing in chronic ER stress is similar to control cells, we examined the subcellular localization of proinsulin during adaptive chronic ER stress and subsequent recovery. Interestingly, an intracellular and cell-periphery localized proinsulin puncta were observed during stress, but not during recovery (**Fig. 1I**). The sequestration of proinsulin during adaptive chronic ER stress into intracellular structures suggests proinsulin processing may be inhibited (Haataja et al., 2013). Taken together, our results show that MIN6 cells develop adaptive translational reprogramming during chronic ER stress with a reversible inhibition of protein synthesis and proinsulin mRNA levels.

### CPA treatment alters the intracellular localization of proinsulin

Due to the essential role of the ER-Golgi transport in insulin maturation, we examined the dynamics of proinsulin cellular localization in MIN6 cells following CPA treatment. In CPA treated cells, proinsulin was highly associated with the ER, shown by the colocalization of proinsulin and P4HB, an ER resident molecular chaperone. CPA treatment for 6h induced a punctate staining of proinsulin in most cells. While the majority of proinsulin was still colocalized with P4HB, the fraction of proinsulin that was localized in puncta did not stain for P4HB. This finding suggested enrichment of proinsulin in granule-like structures outside the ER, near the cell periphery. After 18h of CPA treatment, more than a third of the cells had significant amounts of proinsulin in these peripheral granule-like structures. Interestingly, for these cells, very low proinsulin staining appeared to be colocalized with P4HB (**Fig. 2A-B**), suggesting that proinsulin had largely migrated from the main portion of the ER and became stable in these structures, likely due to decreased processing to insulin.

**Figure 2.**
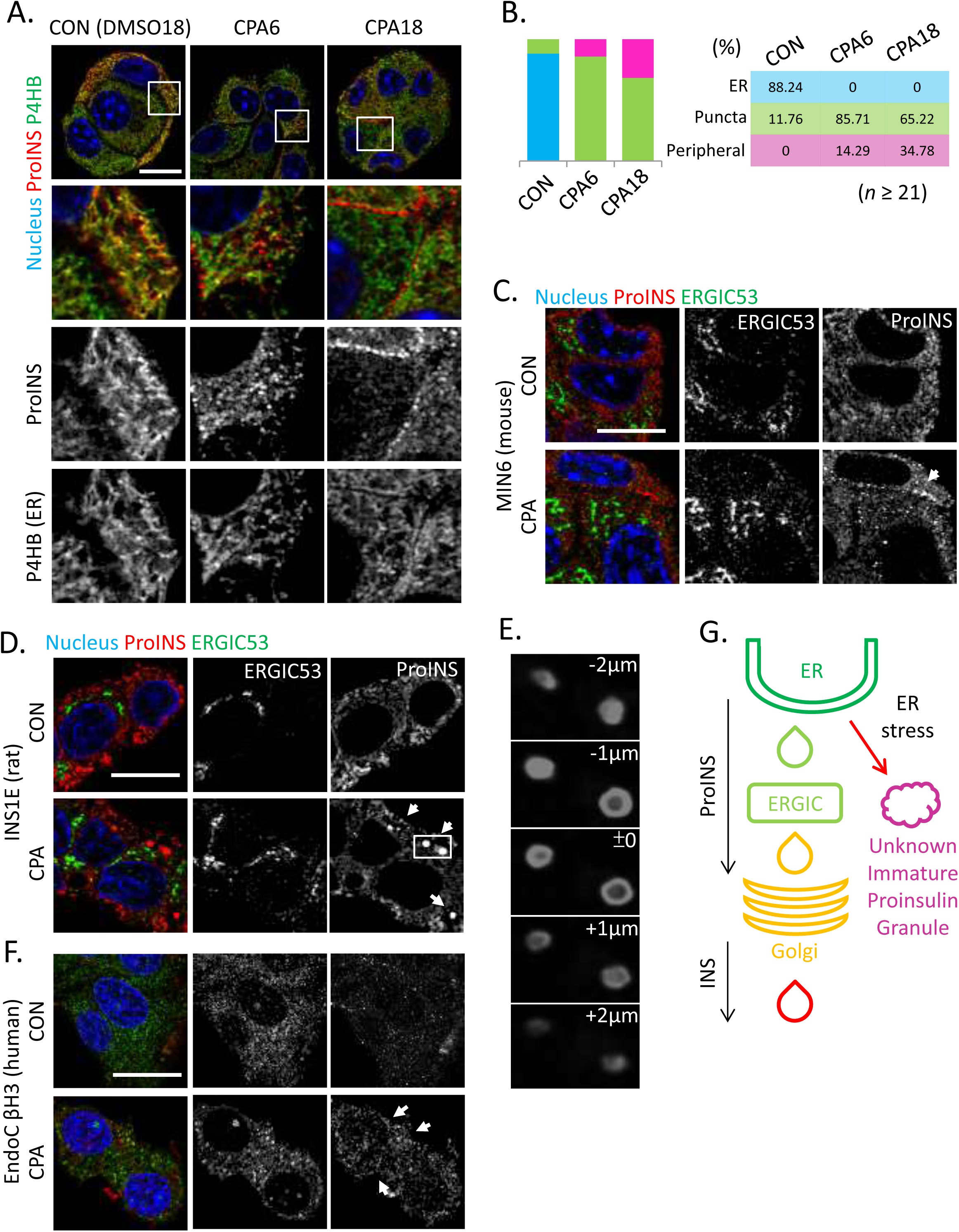
Accumulation of cytoplasmic proinsulin granules in β-cells during chronic ER stress is conserved in rat and human species. (A) Fluorescence immunocytochemistry of MIN6 (A), INS1E (D) and EndoC βH3 (F), cells treated with CPA for 6h and 18h (MIN6), 6h (INS1E) and 72h (EndoC βH3) for the indicated (color-coded) proteins. Boxes indicated the enlarged regions of images below each panel. Scale bar, 10μ. (B) Quantitative analysis of the proinsulin puncta subcellular localization in the indicated treatments of MIN6 cells. (C) Fluorescence immunocytochemistry of MIN6 cells for the indicated proteins and treatments. (E) Reconstruction of stacking images represented by the box in D. (G) Schematic representation of the non-canonical exit of proinsulin from the ER. n, indicates number if cells from 3 independent biological replicates. The % of cells for the indicated phenotype is given in 2B in colored boxes and columns.

We next examined whether proinsulin colocalized with the ER-Golgi intermediate compartment (ERGIC), a transient intracellular structure between ER and Golgi for early protein quality control (Sugawara et al., 2014). We found that there was no colocalization between proinsulin punctate staining and the ERGIC marker ERGIC53 in MIN6 cells in both control and CPA treated conditions (**Fig. 2C**). Finally, we examined whether the punctate CPA-induced proinsulin immunostaining was conserved in β-cells of other species. We examined proinsulin and ERGIC53 in INS1E, a rat insulinoma cell line and EndoC βH3, a human β-cell line, with or without CPA. Large vesicle-like proinsulin-enriched structures appeared following 6h of CPA treatment in INS1E cells (**Fig. 2D and 2E**). The early response of INS1E cells compared to MIN6 cells is consistent with the higher sensitivity of INS1E cells to ER stress. In human EndoC βH3 cells, CPA-induced punctate proinsulin immunostaining was observed after 72h of treatment and did not colocalize with ERGIC53. We observed the proinsulin puncta near the cell periphery in EndoC βH3 cells, which is similar to MIN6 cells (**Fig. 2F**). Collectively, our data suggest that during ER stress adaptation, proinsulin maturation to insulin is impaired and proinsulin accumulates in granule-like structures outside the ER (**Fig. 2G**) whose exact identity remains unknown.

### MIN6 cells undergoing reversible chronic ER stress attenuate expression of genes associated with β-cell identity and function

We have previously shown that ER membrane localized mRNA translation is strongly repressed during adaptive (reversible) chronic ER stress in MEFs (Guan et al., 2017). The high demand of β-cells for ER proteostasis related to the synthesis of proteins critical for β-cell specific functions, motivated us to examine MIN6 cells for β-cell specific changes in gene expression as cells progress from acute to chronic ER stress. We treated MIN6 cells for 1h or 18h corresponding to acute and chronic ER stress (**Sup. Fig. 2A**, CPA1 and CPA18, **Sup. Fig. 2B**). To this end, we analyzed the effects of ER stress on genome-wide perturbations in transcriptomes and translatomes using total RNA sequencing (RNA-seq) in conjunction with ribosome profiling (Chen and Tanaka, 2018; Ingolia et al., 2009). Initially, we analyzed the data for the *Ins2* gene for validation (about 50% of the mRNA in β-cells is contributed by the *Ins* genes). While reading frames of mRNA-seq in *Ins2* mRNA showed no difference, the distribution of ribosome footprints was significantly enriched at reading frame 0 of the mRNA (**Sup. Fig. 2C**), in agreement with ribosome footprints being at the ribosome P site (Chen and Tanaka, 2018; Ingolia et al., 2009). The quality of the data was further confirmed by the distributions of Ribo-seq reads on the *Ins2* mRNA that was selectively enriched within the coding region (**Sup. Fig 2D**), in contrast to the mRNA-seq reads found throughout the gene. In addition, we showed a high correlation among biological replicates for both the transcriptome and translatome datasets (**Sup. Fig. 2E**). Finally, we confirmed that UPR pathway genes were significantly up-regulated in the transcriptome of all three biological replicates (**Sup. Fig. 2F**), further validated by qRT-PCR analysis (**Sup. Fig. 2G**) and Western blotting (**Figure 1A**).

Next, we determined the regulation of the global transcriptome and translatome during acute and chronic ER stress. For the translatomes we used the normalized ribosome footprints to mRNA-seq datasets, and assigned the normalized values the term ribosome occupancy (ribo^ocp^), as an indicator of the translation efficiency of the different mRNAs. We detected in both transcriptomes and translatomes datasets 10,163 transcripts for 6,938 genes. A significant change in mRNA abundance was observed in CPA18, but not CPA1, compared to control cells (**Fig. 3A**), suggesting that reprogramming of the transcriptome is associated with the adaptive chronic ER stress (2036 transcripts were upregulated, while 385 transcripts were downregulated between CPA1 and CPA18). We next assessed differences in ribosome occupancy (ribo^ocp^, Ribo-seq/mRNA-seq) of MIN6 cells treated with CPA (**Fig. 3B**). At CPA18, the total ribo^ocp^ decreased in comparison with the untreated control **(Fig. 3B**), suggesting reprogramming of mRNA translation. We then classified the differentially regulated transcripts into six groups by the trend of changes (up or down) in mRNA abundance (**Fig. 3C**) and ribo^ocp^ (**Fig. 3D**) in response to CPA treatment eliciting acute (CPA1), or chronic (CPA18) ER stress (**Sup. Table 1**). This analysis suggests gene expression reprogramming with progression from acute to chronic ER stress. Although a few mRNAs showed a transient increase or decrease in abundance in the acute response (G2 and G5), the predominant effect was observed in the chronic ER stress with 2036 genes being up-regulated and 385 genes being down-regulated as compared to control cells (G3 and G6). Translational control in the acute response was evident by the 847 mRNAs being ribo^ocp^-up (G8) as compared to 246 mRNAs with continuous up-regulation during progression of stress from acute to chronic conditions (G7). Notably, in agreement with the partial recovery of protein synthesis (**Fig. 1B**), 703 mRNAs decreased their ribo^ocp^ during progression from acute to chronic ER stress (G10). We envision that the chronic ER stress which regulates mRNA abundance and ribo^ocp^ both positively and negatively, favors ER proteostasis and survival. To better understand establishment of the chronic ER stress adaptive proteostasis, we performed pathway analysis of the classified groups, by using the Kyoto Encyclopedia of Genes and Genomes (KEGG) database. Seven groups (G2, 3, 6, 8, 9, 10, 11) showed enrichment of biological functions in the KEGG pathway. Transcripts associated with ribosome biosynthesis, protein processing, degradation and transport, as well as RNA transport and splicing, were significantly upregulated during chronic ER stress. In contrast, functions related to the ribosome, cell cycle and DNA replication were markedly decreased in the acute or chronic stress states, (G10, 11, **Fig. 3E)**. Importantly, transcripts associated with β-cell identity and function (such as *Nkx2-2, Pdx1*, *Glut2*, *Ins1,* and *Ins2*) and function (such as *Glut2*) were substantially decreased between CPA1 and CPA18 (G6, **Fig. 3E**). These results suggest that MIN6 cells can withstand acute ER stress and maintain β-cell specific gene expression. However, β-cell function and identity transcripts are progressively reduced upon establishment of adaptive chronic homeostasis. These data further suggest that loss of β-cells in diabetic islets may be caused by adaptive responses to chronic ER stress without recovery. We will refer to this cellular response as “exhaustive adaptive response” to chronic ER stress, βEAR (<u>β-</u>cell <u>E</u>xhaustive <u>A</u>daptive <u>R</u>esponse).

**Figure 3.**
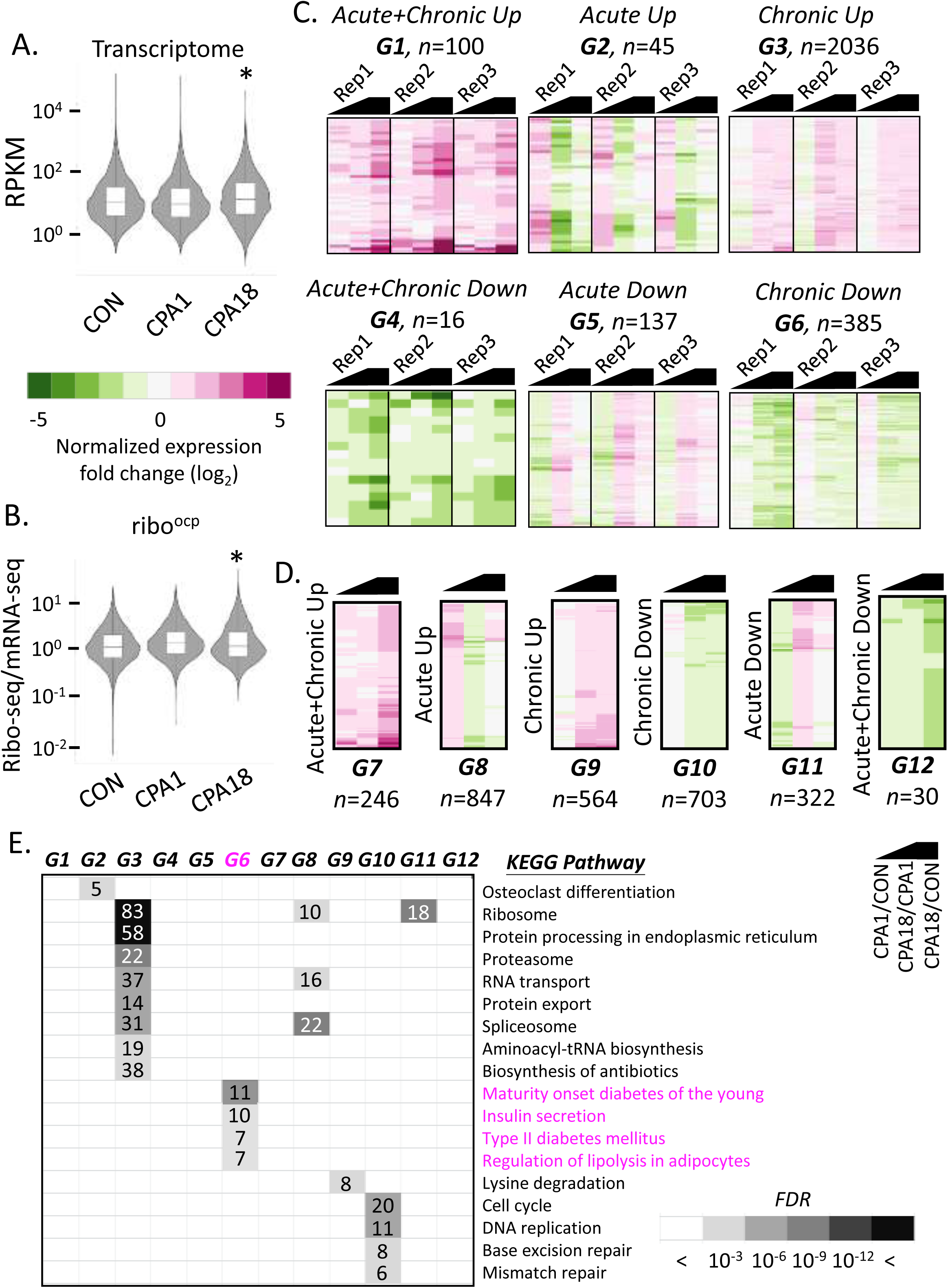
Analysis of transcriptomes and translatomes during progression from acute to chronic phase of ER stress in MIN6 cells. Violin plots representing transcriptomes (A) and translatomes (B), of untreated (CON), 1h CPA treated (CPA1) and 18h CPA treated (CPA18) cells. Heatmaps and the classifications of changes in the expression of transcriptomes (C) and translatomes (D) between CON and CPA1 (CPA1/CON), CPA18 and CPA1 (CPA18/CPA1) and CPA18 and CON (CPA18/CON) datasets. Black symbol indicates the compared datasets. Three biological replicates were used with significance of gene expression, **** p<0.001 Wilcoxon signed-rank test. Letter code G1-G12 indicates the gene sets of assigned common regulation and n, indicates the number of genes per group. (E) KEGG pathway analysis of the classified groups of transcriptomes and translatomes. Boxed numbers indicate the number of genes identified in the individual pathways. *FDR*, False Discovery Rate.

### Transcriptome reprogramming during chronic ER stress involves impaired expression of β-cell related genes

To further assess the temporal changes of the reprogramming (**Fig. 3**) and how they relate to β-cell identity, we performed an additional analysis of the transcriptomes and translatomes of CPA1 vs control (**Fig. 4A**) and CPA18 vs control (**Fig. 4B**), which represented the acute and chronic ER stress phases, respectively (**Sup. Table 1**). In the acute response, we identified 798 mRNAs (11.5%) with increased ribo^ocp^, and 300 mRNAs (4.32%) with a decrease, in agreement with the translational reprogramming during acute ER stress. Changes in mRNA abundance during the acute phase were less prominent, with levels of 138 mRNAs increasing and 108 mRNAs decreasing (**Fig. 4A**). In contrast, there were more changes in mRNA abundance (1795) than translational control (1183) during the chronic response (**Fig. 4B**). Comparing the transition between the acute to chronic phase, it was notable, that changes in mRNA abundance increased from 1.99% (acute) to 20.41% (chronic), while changes in ribo^ocp^ were similar between the two phases. An early increase in mRNA abundance and translation was observed for the *Atf4* gene, that was sustained during the chronic phase, but generally the percent/fold increase in ribo^ocp^ was larger than the percent/fold increase in mRNA levels (**Fig. 4A-B** and **Fig. 1A**). Similarly, downstream of the *Atf4* gene, *Chop* and *Gadd34* showed more than 2-fold increase in mRNA abundance and 3-fold in ribo^ocp^ in the acute phase (**Fig 4A-B**). We also looked at changes in expression of 49 UPR-related genes during the acute and chronic phase (**Fig. 4C** and **Sup. Table 2**). A shift to mRNA abundance was observed for most well-known adaptation genes of the UPR (**Fig. 4C** and **Sup.Table 2**), as well as prosurvival genes, such as transducing β-like 2, *Tbl2* (Tsukumo et al., 2014), stress-associated endoplasmic reticulum protein 1, *Serp1* (Yamaguchi et al., 1999), Wolframin ER Transmembrane Glycoprotein, *Wfs1* (Fonseca et al., 2005) and Selenoprotein S, *SelS* (Qin et al., 2016). In conclusion, β-cells show an enhanced response compared to MEFs (Guan et al., 2017) during the acute phase of ER stress, with elevated mRNA abundance and ribo^ocp^ for important stress adaptation proteins.

**Figure 4.**
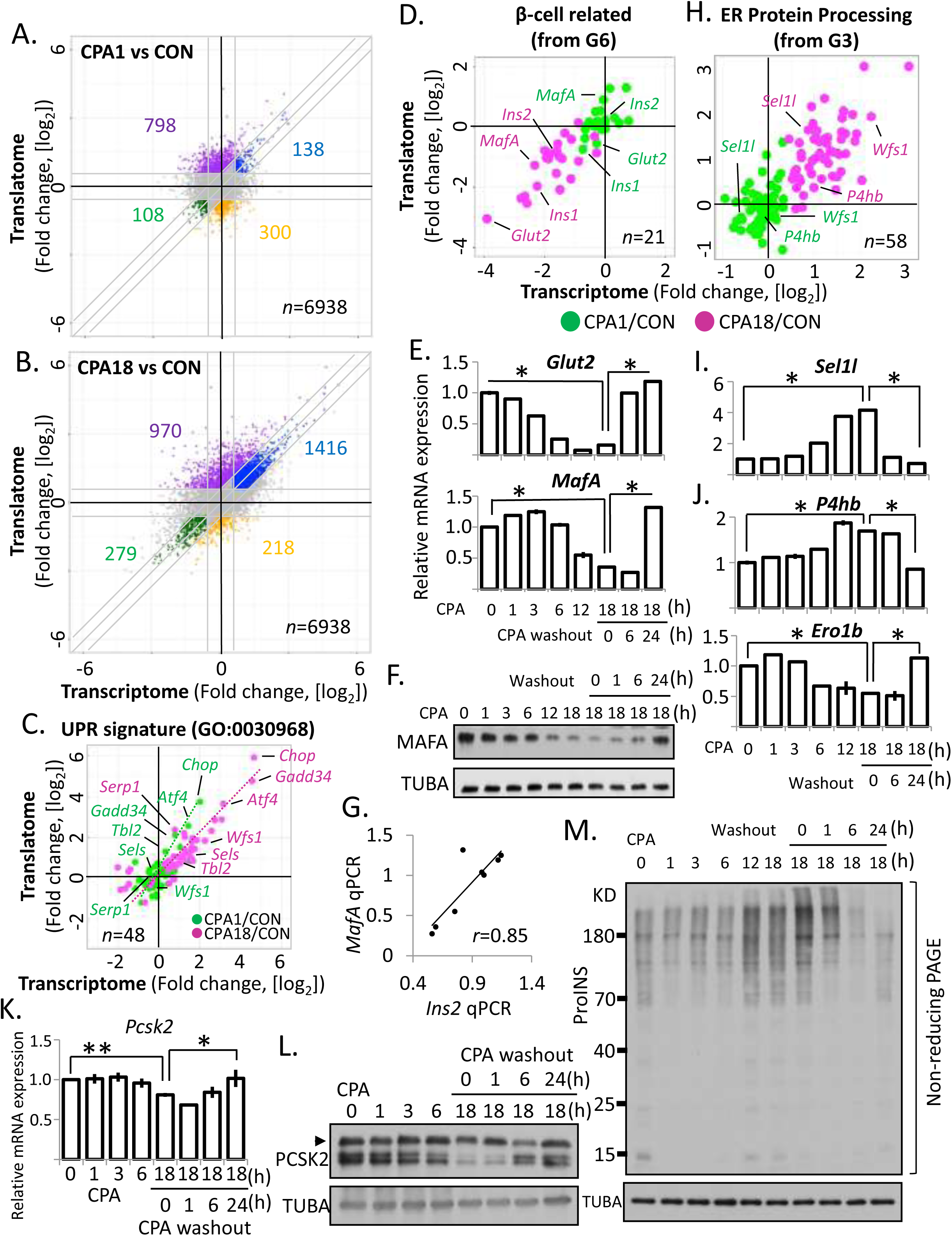
Transcriptional and translational reprogramming in MIN6 cells in response to CPA induced ER stress. Scatterplots of fold changes in CPA1 vs CON (A) and CPA18 vs CON (B). (C and H). Scatterplot of fold changes of the UPR signature (GO:0030968) gene expression between acute (CPA1/CON) and chronic (CPA18/CON) ER stress. (D) Scatterplot of fold changes of genes in groups identified in Fig. 3E (G6 and G3) in acute (CPA1/CON) and chronic (CPA18/CON) ER stress. (E and I) qRT-PCR analysis for the indicated mRNAs normalized to *GAPDH* mRNA levels, in MIN6 cells treated with CPA for the indicated times. (F and L) Western blot analysis for the indicated proteins in MIN6 cells. (G) Scatterplot representing an association between *Ins2* and *MafA* mRNA levels measured by qRT-PCR. (K) qRT-PCR analysis of *Pcsk2* mRNA levels normalized to *GAPDH* mRNA levels, in MIN6 cells under the indicated treatments. (M) Western blot analysis of proinsulin in non-reducing PAGE electrophoresis.

The β-cell specific transcriptome is important to maintain β-cell functions, such as proinsulin biosynthesis and processing, storage and glucose-stimulated insulin secretion (GSIS) (Remedi and Emfinger, 2016). Mis-regulation in one or more regulatory factors in this pathway can result in the pathogenesis of diabetes (Marchetti et al., 2017; Rutter et al., 2015). We found that the mRNA abundance of β-cell-related genes, such as MAF BZIP Transcription Factor A (*MafA*) (Nishimura et al., 2015; Zhang et al., 2005), and *Glut2* (G6 group **Fig. 3E**), a glucose transporter essential for glucose import in rodent β-cells (Guillam et al., 1997; Thorens et al., 2000; Unger, 1991), decreased during the chronic but not acute phase of ER stress (**Fig. 3E** and **Fig. 4D**). Notably, *MafA*, *Ins2* and *Glut2* mRNA abundance, were highly sensitive to prolonged CPA treatment (**Fig. 4D**). Both *Glut2* and *MafA* mRNA levels were severely decreased during chronic ER stress and rapidly increased to pre-treatment levels after CPA washout (**Fig. 4E**). The decrease of *MafA* mRNA levels during chronic ER stress and fast recovery during stress removal, were consistent with the accumulation of MAFA protein levels (**Figure 4F**). Since *MafA* is a transcriptional activator of the insulin gene (Olbrot et al., 2002), we observed the expected positive association between *MafA* mRNA levels and *Ins2* mRNA levels, supporting the idea that the chronic ER stress induced transcriptional reprogramming distinctly inhibits β-cell function (**Fig. 4G**).

The genes associated with ER protein processing were upregulated in chronic ER stress, including molecular chaperones and factors involved in ER associated degradation (ERAD) and ubiquitination (**Fig. 3E** and **Fig. 4H**). For example, we identified an increase of VCP interacting membrane selenoprotein (*Vimp*), Suppressor/Enhancer of Lin-12-like (*Sel1l*) and Derlin-2 (*Derl2*) in chronic ER stress, suggesting an active role for ERAD in the degradation of proteins, including misfolded proinsulin in β-cells (Hoelen et al., 2015). We confirmed that mRNA levels were increased by CPA treatment by qRT-PCR, and after washout, *Sel1l* mRNA levels returned to basal levels (**Fig. 4I**). We noticed that ER molecular chaperones, prolyl-4-hydroxylase (*P4hb*) and endoplasmic reticulum oxidoreductase 1 beta (*Ero1b*) encoding genes were differentially regulated during chronic ER stress and recovery from stress. While *P4hb* mRNA levels were upregulated, *Ero1b* mRNA levels decreased during chronic ER stress (**Fig. 4J**). Furthermore, we found that chronic ER stress, impaired both, mRNA abundance and maturation of the convertase subtilisin/Kexin Type 2 (*Pcsk2*), a peptidase essential for insulin maturation (**Fig. 4K-I**). Because molecular chaperones are important in proinsulin disulfide bond formation and maturation, we tested proinsulin misfolding in β-cells during ER stress. We used non-reducing PAGE gel electrophoresis and found a gradual increase of high molecular weight disulfide-linked proinsulin complexes upon CPA treatment. CPA washout was sufficient to rescue this phenotype over time (**Fig. 4M**). Collectively, the balanced regulation of ER quality control genes resulted in reversible ER stress, however, it also compromised proinsulin gene expression and protein maturation.

### ER-Golgi transport plays a major role in adaptation to ER stress in MIN6 cells

It is currently unknown whether secretory cells utilize a specific cellular machinery to achieve adaptive homeostasis as compared to non-secretory cells. Thus, we compared the transcriptomes of the acute ER stress and chronic ER stress adaptation in MIN6 and MEFs. This analysis in MIN6 cells identified 1534 mRNAs with increased abundance and 307 mRNAs with decreased abundance, between the acute and the adaptive timepoints (**Sup. Fig. 3A**). Similarly, we identified 603 and 296 mRNAs subjected to positive or negative translational control, respectively. mRNAs encoding for proteins of the ribosome biosynthesis pathway were enriched in the 603 group, while the ribo^ocp^ of cell cycle related mRNAs was reduced (**Sup. Fig. 3A**). Re-analysis of pre-existing sequencing data from MEFs (Guan et al., 2017) identified 567 and 1247 mRNAs which showed upregulation or downregulation of mRNA abundance (**Supp. Fig. 3A**). KEGG pathway analysis of the down regulated genes between MIN6 and MEFs, showed the stress responses were cell type specific, and tailored to cellular function: a decrease of β-cell related functions was observed in MIN6 cells, while mRNAs of cell adhesion and ECM were decreased in MEFs during the transitioning from acute to chronic stress adaptation (**Supp. Fig. 3B**).

In corroboration with common adaptation mechanisms reported among different cell types (Kultz, 2003), in the subset of mRNAs with increased abundance, we identified an enrichment of genes in the ER protein processing pathway (**Sup. Fig. 3C**). 68 mRNAs showed increased abundance and 10 mRNAs showed translational upregulation of this cellular pathway in MIN6 cells. In MEFs, a smaller proportion of mRNAs (37 of 78), were regulated, suggesting similar, less elaborate adaptation machinery in the non-secretory cells. We analyzed the 37 overlapping genes between MIN6 and MEFs by Gene Ontology (GO) in Biological Process (Mi et al., 2019). We observed that these genes were highly associated with ER stress amelioration, including molecular chaperones and ERAD. The 41 MIN6-specific regulated genes were enriched for functions in ER-to-Golgi vesicle transport (**Sup. Fig. 3D**). This analysis revealed a unique adaptive event in MIN6 cells, via an enhancement of the first step of vesicular transport, suggesting that the stress adaptation is influenced by cellular function. Thus, we term this group of 78 genes in MIN6 cells, the β-cell-specific adaptosome.

### Lysosome function is impaired during chronic ER stress

Accumulation of misfolded proteins could be a consequence of proteostatic imbalance (Hipp et al., 2014). Because intracellular immature proinsulin granule-like structures form, despite active ERAD (Hou et al., 2009), we hypothesized that a novel mechanism may be involved. Formation of proinsulin granules was reversible, supporting their formation as an adaptive response to chronic ER stress (**Fig. 1I**). To shed light on the mechanism of proinsulin accumulation, we examined gene regulation (up- or down) in chronic ER stress (**Fig. 5A**). We noted that subunits of the proteasome were upregulated (**Fig. 5B**). By pathway analysis of the down-regulated 497 mRNAs (**Fig. 3B**, combined 279 and 218 mRNAs), we identified lysosome biogenesis genes as significantly repressed (**Fig. 5A**), such as, *Arsb*, a lysosomal arylsulfatase, and *Lamp2*, a lysosomal associated membrane protein (**Fig. 5C**), suggesting impaired lysosome function in chronic ER stress. Accordingly, we examined whether the lysosomal bioactivity was affected. We observed a time dependent reduction of lysosomal enzymatic activity during CPA treatment. MIN6 cells treated with Bafilomycin A1 (Baf A1), an inhibitor of the lysosomal proton pump V-ATPase that serves as a positive control, for 4h caused a 50% reduction of lysosome activity compared to untreated control. In parallel, 18h of CPA exposure showed a 45% reduction of the lysosome activity, which gradually returned to the untreated state 6h after CPA washout (**Fig. 5D**). We monitored the protein level of the autophagy substrate ubiquitin-binding protein p62, encoded by the sequestrosome 1 gene, *Sqstm1*, and the microtubule-associated protein 1A/1B-light chain 3, LC3-I to LC3-II conversion in MIN6 cells during CPA treatment and after washout. We observed a gradual increase of p62 protein and LC3-II levels during CPA treatment, which was similar to Baf A1-induced interruption of autophagosome degradation by the lysosome (**Fig. 5E**). Lastly, we observed an increase of punctate LC3 staining in MIN6 cells after 18h of CPA treatment (**Fig. 5F**). Taken together, we conclude that the pathway of autophagosome-to-lysosomal degradation was inhibited during chronic ER stress in MIN6 cells.

**Figure 5.**
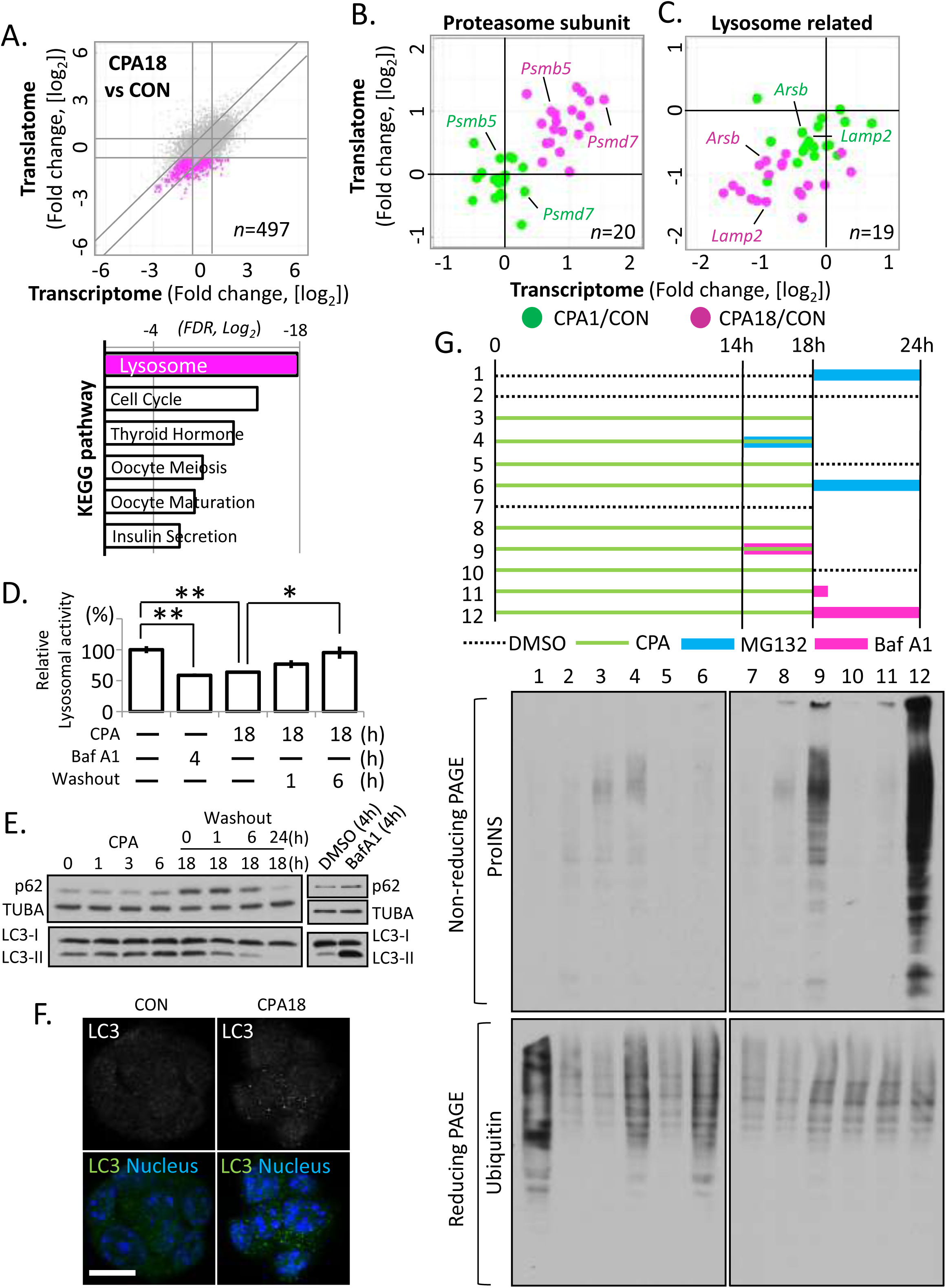
Impairment of lysosome activity correlates with misfolded proinsulin accumulation during chronic ER stress. (A) Scatterplot of fold changes in CPA18 vs CON MIN6 cells, with the repressed gene expression indicated in pink, and the KEGG pathway analysis of the highlighted group, below panel A. (B-C) Scatterplots of fold change in acute (CPA1/CON) and chronic (CPA18/CON) ER stress for genes related to the proteasome (B) and the lysosome (C). (D) lysosome enzymatic activity measured in MIN6 cells treated with Baf A1 or CPA. (E) Western blot analysis of the indicated proteins in MIN6 cells. (F) Fluorescence immunocytochemistry for LC3, in MIN6 cells treated with CPA for 18h. Nuclei are indicated with staining in blue. Scale bar is 20μ. (G) *upper*, schematic representation of 12 different MIN6 treatments with CPA, MG132 or Baf A1. Green lines indicate CPA treatment. CPA was washed out between 18-24h; *lower*, Western blot analysis for proinsulin by non-reducing PAGE electrophoresis in extracts prepared from the 12 MIN6 treatments. Western blot analysis for ubiquitin by reducing PAGE electrophoresis.

### Impairment of lysosome activity correlates with misfolded proinsulin accumulation

The ubiquitin-proteasome system (UPS) and the lysosome are the two major quality control systems for proteostasis. Our analysis demonstrated that during chronic ER stress in MIN6 cells, lysosomal activity gradually declined, while the molecular pathway associated with UPS was up-regulated (**Fig. 3E**, **Fig. 4H**, **5C**). Moreover, our analysis revealed an accumulation of misfolded proinsulin outside the ER upon CPA treatment (**Fig. 2** and **Fig. 4M**), implying disrupted proinsulin proteostasis in chronic ER stress. Accordingly, we determined if the proteostatic pathways, UPS or lysosome, were involved in this regulation. We treated MIN6 cells with CPA for 18h together with MG132, a proteasome inhibitor, or Baf A1 and monitored the proinsulin folding status by non-reducing PAGE electrophoresis and western blotting. Lysosomal impairment by Baf A1 treatment over the last 4h of the 18h of CPA treatment strongly increased the amount of misfolded proinsulin (**Fig. 5G**, lane 9). Moreover, a 6h CPA washout in the presence of Baf A1 led to a dramatic accumulation of misfolded proinsulin (**Fig. 5G**, lane 12), while MG132 caused a modest effect. Therefore, our data suggest that effects of CPA-mediated chronic ER stress on autophagosome-to-lysosomal degradation impairs clearance of misfolded proinsulin.

### Chronic ER stress adaptation genes identified in MIN6 cells are downregulated in type 1 diabetes

Pancreatic β-cells are long-lived; therefore they encounter numerous changes in their microenvironment over their life span, such as the daily feeding-fasting cycles. Hyperglycemia, hyperlipidemia and starvation, are all inducers of ER stress (Fonseca et al., 2011). Indeed, several studies have highlighted the importance of the ER stress response in diabetes mellitus (Eizirik et al., 2008; Ozcan et al., 2004). We found an accumulation of misfolded proinsulin and the formation of immature proinsulin in granule-like structures during the transition from acute to chronic ER stress adaptation (**Fig. 2** and **Fig. 4M**). In addition, the chronic ER stress induced transcriptional reprogramming, largely suppressed expression of β-cell specific genes (**Fig. 4D** and **Sup. Fig. 3**). Therefore, we hypothesized that β-cell loss/dysfunction in diabetes could be associated with the homeostatic state of ER stress adaptation or the failure of it.

To test this hypothesis, we selected genes that were affected in MIN6 cells during the chronic adaptive state, and filtered out those that exhibited similar behavior in MEFs (Guan et al., 2017). This allowed us to establish a β cell-specific ER stress gene set. This gene set, which contains 334 genes was enriched for β-cell associated functions by the KEGG pathway analysis and we called it regulatome (**Fig. 6A**). We examined the expression levels of this gene set in a pre-existing microarray database (Holm et al., 2018; Trembath et al., 2020) and the sc-RNAseq data produced by the Human. Pancreas Analysis Program (HPAP) accessible in PancDB (Kaestner et al., 2019). The microarray data were generated from laser-excised islets from patients with T1D and T2D and healthy donors (**Fig. 6B**). We identified 11 genes that showed downregulation in T1D samples, compared to healthy controls (**Fig. 6B**), but not in T2D samples (**Sup. Fig. 4A**). One gene, *PPP1R1A*, was identified in both T1D and T2D, but showed less significant change in T2D. 10 out of the 11 downregulated genes were present and decreased in MIN6 cells during chronic ER stress adaptation (**Fig. 6B**).

**Figure 6.**
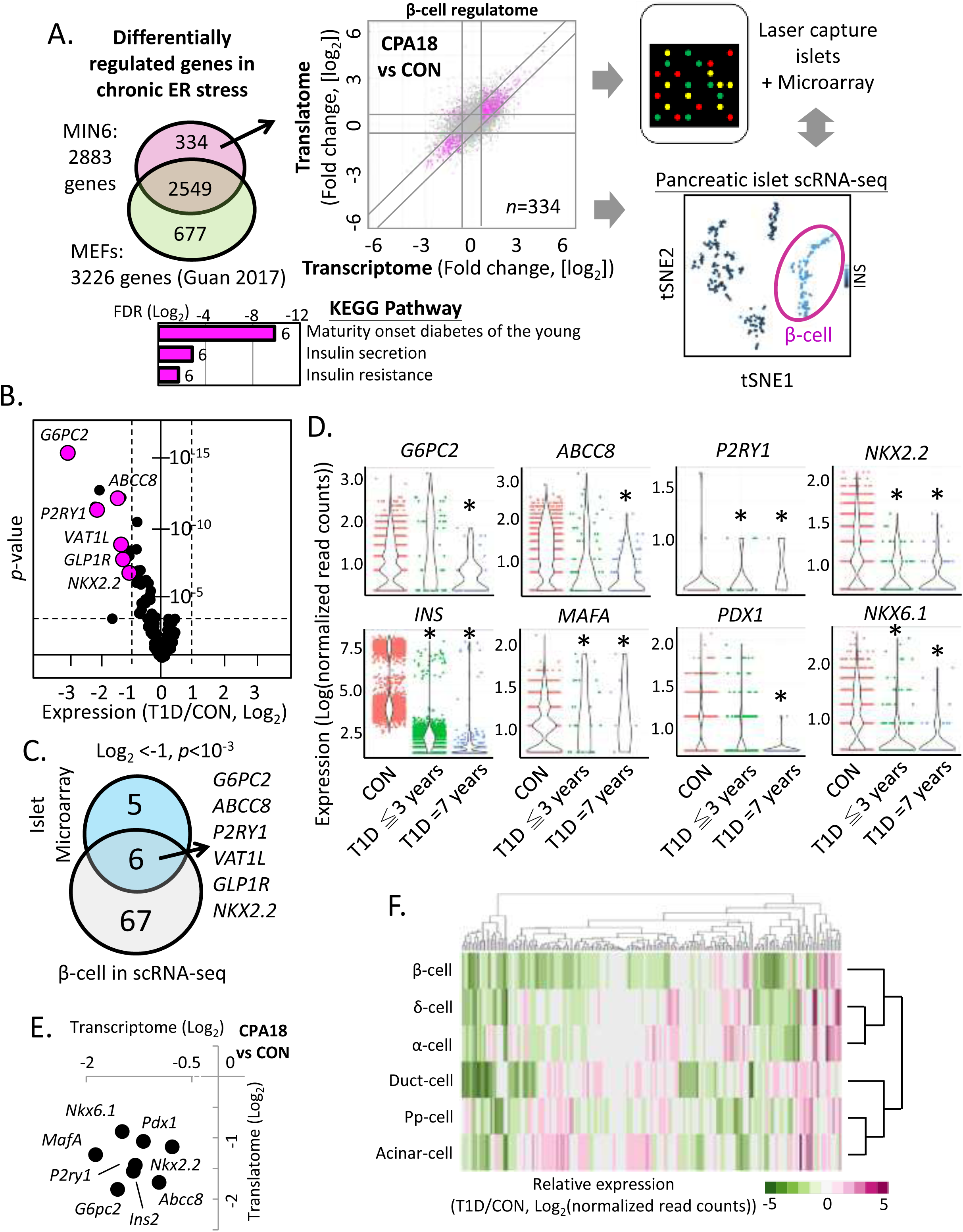
Altered expression of the β-cell ER stress-induced gene set (regulatome) in T1D islets reveals biomarkers of T1D progression. (A) Schematic representation of the β-cell ER stress gene set (334 genes) in MIN6 cells and determination of its expression in T1D islets or scRNA-seq datasets. Scatterplot indicates in color the expression patterns of the 334 gene set during chronic ER stress in MIN6 cells. The KEEG pathway of the 334 gene set is shown below panel A. (B) The 334 gene set expression profile in T1D islet microarray datasets, shown by a volcano plot. (C) Venn-diagram showing the common repressed genes of the 334 gene set in T1D, between islet microarray and β-cells scRNA-seq datasets. (D) Expression of genes in β-cells scRNA-seq datasets of healthy donors (CON), early T1D (T1D ≦ 3 years) and prolonged T1D (T1D = 7 years) patients. (E) Scatterplot of fold change in MIN6 cells during chronic ER stress (CPA18 vs CON). (F) Heatmap of gene expression profiles of the 334 gene set in different cell types of T1D scRNA-seq datasets, compared to healthy donors.

Pancreatic islets, known as the islets of Langerhans, consist of several cell types. Normal human islets have α-cells (30-50%), β-cells (50-60%), δ-cells (<10%), pp-cells (<5%) and ε-cells (<1%). In T1D patients, there is a specific loss of β-cells (Campbell and Newgard, 2021; Katsarou et al., 2017), thus the 11 down regulated genes in the T1D pancreatic islets microarray dataset could potentially be caused by alternative islet cell types. To further investigate β-cell specific responses in T1D, we used the scRNA-seq from HPAP that analyzed islets of T1D patients and healthy donors (Kaestner et al., 2019). We first compared the expression profiles of healthy donors and T1D patient islets for the 11 genes in the scRNA-seq dataset (**Sup. Fig. 4B**). We found that the expression of the 11 genes was decreased in T1D β-cells (**Sup. Fig. 4C**), so we applied a more stringent cut off (FDR≤10^-3^) and observed 6 of the 11 genes were significantly downregulated in β-cells of T1D patients (**Fig. 6C-D**). In addition to the 11 genes, *INS* and *MAFA* mRNA abundance was also decreased in β-cells of patients with T1D (**Fig. 6D**). In T1D β-cells, *G6PC2*, *ABCC8* and *NKX2.2* showed a significant decrease in expression in comparison to healthy donors (**Fig. 6C**). All 8 genes shown in Fig. 6D, were also down-regulated in MIN6 cells during chronic ER stress (**Fig. 6E**). Lastly, we examined how the expression of the regulatome gene set (**Fig. 6A**) is regulated in the scRNAseq dataset of T1D islets (**Fig. 6F**). We observed a higher decrease in expression of these genes in β-cells in comparison to other cell types in T1D (β-cells/ all other cell types = 57.23%: 29.13 ± 0.79%). Only 7.51% of the gene set showed an increase in expression in β-cells, which was lower than other cell types (12.95 ± 2.23%) (**Fig. 6F**). Collectively, our analysis revealed reduced abundance of distinct mRNAs of the β-cell chronic ER stress gene set in T1D (**Fig. 6A**). This finding supports the hypothesis that there is likely a failure of chronic ER stress adaptation in β-cells in T1D that leads to targeted downregulation of β-cell function and identity-specific mRNAs. We propose that failure to adapt to chronic ER stress, or as we called in this report, βEAR may contribute to the pathogenesis of T1D in patients.

### Frequent episodes of chronic ER stress in MIN6 cells lead to decreased plasticity and gradual loss of β-cell specific gene expression

The decreased expression of β cell specific genes in T1D islets and β-cells from T1D patient islets (**Fig. 6B-D**), suggested inadequate adaptation to chronic ER stress in β-cells. To test this hypothesis, we compared the expression profile of the MIN6 β-cell-specific adaptosome (78 genes) with the scRNA-seq data from the combined T1D and healthy donor β-cells for similarities in gene expression. We found overlap of 46 out of the 78 genes which were expressed in both healthy donors or T1D patients. As opposed to upregulation during the chronic adaptation state in MIN6 cells (**Sup. Fig. 3)**, most of the adaptosome genes were downregulated in T1D patients compared to healthy donors (**Fig.7A**). Notably, the expression of *BiP*, also known as *HSPA5*, was significantly increased in T1D patients (**Fig.7A**). *BiP* is a key modulator of the UPR (Wang et al., 2009) and is considered a UPR sensor (Kopp et al., 2019). Increased abundance of *BiP* mRNA in T1D β-cells without a similar increase of the ER adaptosome genes, suggests the presence of an unresolved UPR. These data reveal the sustained loss of induction of the adaptosome in β-cells from T1D patients under conditions of unresolved ER stress.

**Figure 7.**
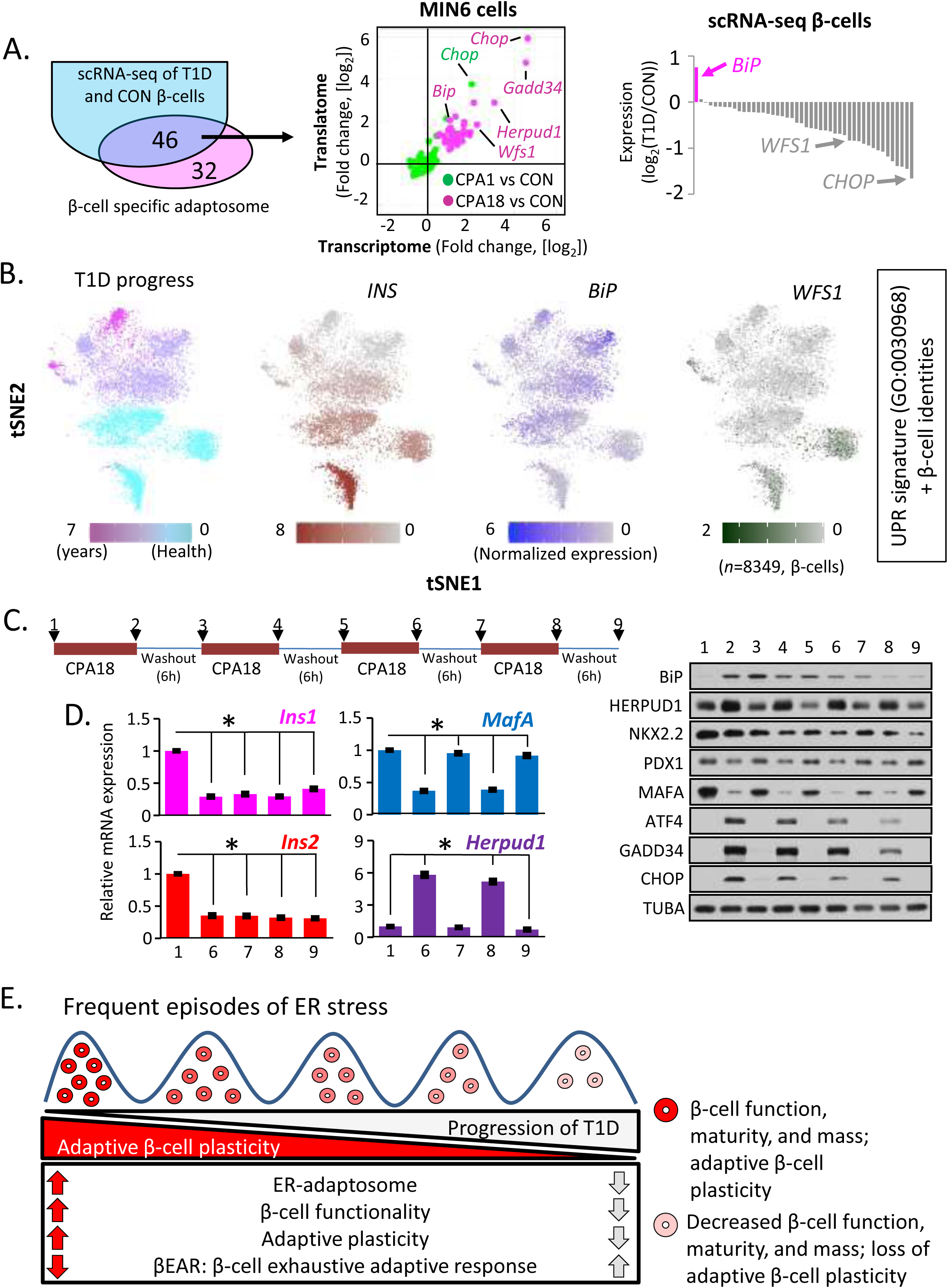
Progression of T1D pathogenesis correlates with ER-stress-induced adaptive exhaustion and suppression of β-cell maturity markers. (A) Schematic representation of the common genes between the β -cell specific adaptosome in MIN6 cells (78 genes) and the β-cells scRNA-seq datasets. Plots show the regulation of the 46 genes in MIN6 cells during acute and chronic ER stress (middle) and β-cells scRNA-seq from healthy and T1D patients (right). (B) Relevant distance in gene expression between two data sets: (tSNE1) scRNA-seq dataset from all human β-cells (healthy and T1D, 8,349) and (tSNE2) the expression of a group of genes consisting of both, the UPR signature genes (GO:0030968), and 6 β-cell-specific gene markers (INS, PAX6, NKX2.2, NKX6.1, MAFA and PDX1). Colored from left to right, T1D progression, *INS*, *BiP* and *WFS1*. Darker color indicates higher expression of the respective genes. Raw data were normalized as described in Materials and Methods. (C) Schematic of stress/recovery cycles and Western blot analysis of UPR and β identity genes. (D) qRT-PCR analysis of RNA levels normalized to *GAPDH* mRNA, for *Ins1*, *Ins2*, *MafA* and *Herpud1* mRNAs in the last 2 stress/recovery cycles. *n*=3 biological replicates. Error bars represent S.E.M. *. p<0.01 two-tailed paired Student’s t-test. All comparisons were to sample 1. (E) Working hypothesis model of β EAR in T1D pathogenesis.

Next, we investigate looked at the relevant distance in gene expression between two data sets: (i) scRNA-seq dataset from all human β-cells (healthy and T1D, 8,349) and (ii) the expression of a group of genes consisting of both, the UPR signature genes (GO:0030968, (You et al., 2021)), and 6 β-cell-specific gene markers (*INS, PAX6, NKX2.2, NKX6.1, MAFA* and *PDX1)* (**Fig. 7B**, x and y axis, respectively). The similarity metric was determined via the t-distributed stochastic neighbor embedding algorithm (van der Maaten and Hinton, 2008), t-SNE, between the above two data sets, tSNE1 and tSNE2 (**Fig. 7B**). This analysis identified clusters of β-cells with the UPR signature but heterogeneous UPR marker gene expression (**Fig. 7B**). First, by mapping the T1D disease progression (**Supp. Fig 4B**) on the UPR signature clusters of β-cells, we identified clusters corresponding to healthy donors and others derived from patients with more advanced T1D (**Fig 7B**). Second, we determined the expression of *INS* mRNA in each cluster, to define β-cell identity, and found a good correlation between high levels of *INS* gene expression and the healthy donors, as well as noticed a decline of *INS* mRNA abundance with T1D pathogenesis (**Fig. 7B**). Furthermore, we observed an anti-correlation between *INS* and *BiP* gene expression with T1D pathogenesis (**Fig. 7B**), indicating an unresolved ER stress response with progression of T1D. In agreement with unresolved ER stress (**Fig. 7A**), the β-cell ER proteostasis adaptosome marker *WFS1*, was decreased in T1D pathogenesis (**Fig. 7A**), and was more abundant in the healthy β-cell clusters. Taken together, these data suggest that β-cells from T1D patients experience an unresolved UPR.

It has been recently reported that preserving heterogeneity in β-cell maturity in islets is essential for their function *in vivo* (Nasteska et al., 2021). Thus, our data showing that ER stress decreases expression of genes involved in β-cell maturity, combined with the documented heterogeneity of β-cells in islets *in vivo*, suggests that β-cells undergo cycles of ER stress and recovery in response to metabolic stress. We therefore reasoned that the collapsing chronic ER stress adaptation response in T1D islets could result from exposure to frequent ER stress cycles that fail to return to homeostasis. Alternatively, β-cells may fail to recover after prolonged ER stress. We therefore tested these two hypothesis in MIN6 cells. We first treated MIN6 cells with 6, 18 or 42h of CPA, followed by a 24h of recovery (**Sup. Fig.5A**). The UPR markers ATF4, GADD34, CHOP and BiP, showed complete reversibility after prolonged stress (**Sup.Fig. 5B**). Although prolonged ER stress decreased cell survival (**Sup.Fig. 5C**), the surviving cells recovered after stress washout (**Supp. Fig. 5B**). Our data suggest that β-cells can endure prolonged ER stress and recover from it.

We next determined the effect of cycles of 6h recovery time (compare **Fig. 7C** with **Supp. Fig. 5A**) from 18h chronic ER stress exposure on the levels of stress-induced and β-cell specific proteins. MIN6 cells were challenged with four cycles of 18h of CPA treatment then followed by a 6h of recovery in fresh medium (**Fig. 7C**). We examined the protein expressions of both UPR genes and β-cell identity markers in response to the stress/recovery cycling. We found a gradual reduction of UPR proteins (ATF4, CHOP, GADD34 and HERPUD1) with increasing cycle number. The expression of *BiP* was induced in the first cycle of CPA treatment, but increased expression was not observed in subsequent cycles (**Fig. 7C**). Although these data suggest *BiP* regulation is lost with stress/recover cycling, the levels of the protein remained higher in cycling than untreated cells (**Fig. 7C**, compare lanes 1 with 5, 7 and 9). Notably, the β-cell marker, NKX2.2, showed a decrease and partial recovery during the second and the third stress/recovery cycle, but was unchanged following the first stress cycle. NKX2.2 protein levels gradually decreased with cycles of stress (**Fig. 7C**). MAFA, on the other hand, was affected by early stress cycles; MAFA showed a significant decrease, with partial recovery, at the end of the first and subsequent cycles (**Fig. 7C**). Lastly, we examined the effects of stress cycles in *Ins1*, and *Ins2* mRNA levels by qRT-PCR during the last two stress cycles. A significant irreversible reduction in *Ins1* and *Ins2* gene expression was observed (**Fig. 7D**). The pattern of changes in *MafA* and *Herpud1* mRNA levels in the last two cycles were in good correlation with the pattern of changes in their protein levels (Fig. 7D and 7C). Taken together, these results show increasing frequency of stress/recovery cycling progressively diminish β-cell plasticity and return to homeostasis. Since T1D is a condition of increased β-cells sensitivity to environmental and nutritional factors, we propose that frequent episodes of chronic ER stress on β-cells may contribute to diminution of β-cell identity and promote β-cell loss (**Fig. 7E**).

## Discussion

T1D is preceded by β-cell dysfunction in islets, but the mechanisms of this response are not well understood (Rothman and Greenland, 2005; Tersey et al., 2012). The presence of ER stress markers in β-cells of mouse models and human islets of T1D patients (Engin et al., 2013; Marhfour et al., 2012; Marre et al., 2015; Tersey et al., 2012), indicates that ER stress could cause β-cell dysfunction and progression of T1D (Engin, 2016). In the T1D microenvironment, ER stress can develop by several factors, inherent to β-cells or in response to cytokines secreted by local T cells and macrophages (Tersey et al., 2012). Here, we tested the intrinsic effect of ER stress and the impact of recovery from it on mature β-cell identity as a means to understand the pathogenesis of T1D. Using a reversible chemical that elicits a bona fide ER stress response, we revealed hidden plasticity of β-cells during their response to chronic ER stress. We showed that during chronic ER stress, β-cells undergo a homeostatic state change associated with pro-survival reprogramming of their transcriptome and translatome. Chronic β-cell ER stress altered more than half of the genes known to participate in ER protein processing while at least temporarily compromising β-cell identity. Upon relief from stress, β-cells could regain their mature identity, indicating plasticity. This suggests that episodes of ER stress, even at high amplitude, are tolerated by β-cells in a manner designed to protect ER function.

Importantly, frequent repeated cycles of ER stress diminished β-cell plasticity, as shown an inability to recover their identity genes, the most prominent being *Ins1* and *Ins2*. Laser captured islets and single-cell RNAseq data from patients with T1D (HPAP), indicated gene expression alterations in T1D β-cells, consistent with a model of ER stress adaptive exhaustion and deregulation of the β-cell ER adaptosome during disease progression. It thus appears that frequent cycles of ER stress over time exhaust the adaptation machinery of β cells to promote an ultimately irreversible loss of function that likely facilitates T1D pathogenesis.

We have previously shown that non-secretory cells survive chronic ER stress by reprogramming their mechanism of mRNA translation (Guan et al., 2017). The most efficient mechanism of mRNA translation initiation is the recruitment of the eIF4F complex to the 5’-end mRNA cap, which in turns assembles the preinitiation complex with the 40S ribosome, the multi-subunit protein eIF3 and the Met-tRNA_i_/ternary complex (Merrick, 2015; Pelletier and Sonenberg, 2019). In normal cells, almost all mRNAs are translated via this mechanism. However, cells inactivate this mechanism during chronic ER stress, and inhibit mRNA translation in a manner dependent on activation of the PERK kinase and eIF2α phosphorylation (Costa-Mattioli and Walter, 2020; Guan et al., 2017). Next, selective mRNA translation encoding for proteins promoting adaptation is activated via direct recruitment of the ribosome to these mRNAs by the translation initiation factor eIF3 (Guan et al., 2017). This translational reprogramming significantly dampens ER-associated mRNA translation, while maintaining translation of important transcriptionally-induced mRNAs for the adaptation process. Therefore, the adaptive translational reprogramming is a pro-survival mechanism that decreases ER load and balances ER protein processing to maximize ER function and protein secretion during ER stress (Hakonen et al., 2018; Meyer and Doroudgar, 2020).

How do β-cells adapt to ER stress? Considering the abundance of the *Ins* mRNA translated on ER membranes, ER stress should impair proinsulin synthesis, unless the proinsulin mRNA is part of the selective adaptive translation initiation program. However, if there is selective proinsulin mRNA translation during chronic ER stress, how would β-cells handle the load of making large amounts of proinsulin on the ER? We discovered that β-cell adaptation to chronic ER stress involves dramatic decreases in *Ins* mRNA levels accompanied by dynamic changes in proinsulin protein levels (**Fig. 1H**). The regulatory mechanisms of *Ins* mRNA translation have not been studied here; however, we can speculate that splice-variants of the proinsulin mRNA 5’-UTRs could be a possible mechanism for this regulation. In fact, a splice variant was identified which promotes efficient translation of the *Ins* mRNA in response to metabolic changes in human pancreatic islets (Shalev et al., 2002). Recently, it was shown that the balance between the canonical eIF4E-dependent to the non-canonical eIF3-independent translation initiation is important for cellular adaptation to the availability of glucose (Lamper et al., 2020). These processes can be investigated as potential regulators of *Ins* mRNA translation during chronic ER stress.

A new finding here is that β-cells exposed to chronic ER stress sustain adaptive plasticity. Although chronic ER stress decreases expression of β-cell identity genes, such as *Ins1*, *Ins2*, *Pdx1* and *MafA*, removal of stress restores their expression. The physiological significance of this regulation is likely to decrease *Ins* gene expression transiently, given that PDX1 and MAFA are transcription factors that regulate *Ins* gene expression (Zhu et al., 2017). Plasticity was also observed for the stress-induced genes which returned to the basal levels of expression upon removal of stress. In contrast, frequent repeated ER stress episodes decrease adaptive plasticity for the β-cell identity genes and a subset of the stress-induced genes (**Fig. 7E**). For example, both *Ins1* and *Ins2* mRNA levels recovered after a single cycle of ER stress (**Fig. 1G**) but declined further and did not recover following 3 or more cycles of ER stress and recovery (**Fig. 7D**). The exact mechanisms of adaptive plasticity and exhaustion remain unknown, and considering that translational, transcriptional and mRNA decay programs can all contribute to these cellular responses (Eizirik and Cnop, 2010), only a systematic approach of the effects of various ER-stress signaling pathways (Marre et al., 2015), can reveal the mechanistic details of this regulation.

Although we focused on molecular adaptive plasticity here, metabolic plasticity is also expected to be part of the adaptation mechanism to chronic ER stress in a manner dependent on the cell-type specific functions (Gao et al., 2020; Yadav et al., 2017). To demonstrate that adaptation to chronic ER stress also involves adaptive metabolic plasticity, we determined how cholesterol biosynthesis is regulated, owing to its importance for GSIS (Xia et al., 2008). We found that biosynthesis of cholesterol decreased at chronic ER stress and gradually recovered upon stress removal (**Sup. Fig.6**); the inhibition of cholesterol biosynthesis was consistent with the adaptive response of β-cells to chronic ER stress. While *Ins* mRNA levels decrease, mechanisms of inhibition of secretion of proinsulin or insulin, such as decreased cholesterol biosynthesis (Xia et al., 2008), can generate a reserve pool of insulin for recovery from stress.

Our findings on adaptive plasticity to chronic ER stress are reminiscent of recent studies on β-cell heterogeneity *in vivo*, as a requirement for islet function (Farack et al., 2019; Nasteska et al., 2021). It was shown in a mouse model that adult islets are composed of mature and immature β-cells, determined by the level of expression of *MafA* and *Pdx1*, and the balanced expression of these β-cells populations is important for the physiological metabolic response of the islets (Nasteska et al., 2021). The authors indicated that β-cells isolation from islets can eliminate the differences in maturity, further supporting the plasticity of β-cells in the islet microenvironment (Nasteska et al., 2021). We can speculate that subpopulations of β-cells in the islets, represent cells adapting to ER stress (low levels of *MafA* and *Pdx1*) and cells recovered from ER stress state (high levels of *MafA* and *Pdx1*). Therefore, the heterogeneity in β-cells maturity in normal islets may be a physiological response to cycles of ER stress and recovery, *in vivo*.

How T1D pathogenesis relates to βEAR? Exhaustive adaptation to ER stress could reflect inadequate expression of genes involved in protective adaptive responses and/or permanent decrease of β-cell identity genes, including *INS* gene expression. We showed that the expression of the protective ER adaptosome genes decreased in T1D β-cells (**Fig. 7A**). Only *BiP* mRNA levels increased in T1D β-cells consistent with a state of unresolved ER stress (Kopp et al., 2019). In support of this conclusion, the levels of the *INS* plus 11 additional mRNAs related to β-cell function were decreased. Although we showed, that decreased levels of these genes is part of the adaptive response to chronic ER stress in MIN6 cells (**Fig. 7A**), the permanent decrease accompanying failure of the ER adaptosome response in T1D β-cells, demonstrates a state of failing recovery from stress. We therefore propose as loss of β-cell plasticity in response to frequent cycles of stress in the islet microenvironment as a contributor to T1D pathogenesis, which may involve the loss of balance between mature and immature islet β-cells (Nasteska et al., 2021). The survival of immature β-cells in T1D patients may also associate to their increased proliferation under low insulin levels (Szabat et al., 2016).

Is the positive regulation of the ER proinsulin protein exit an adaptive event in response to chronic ER stress in β-cells? During chronic ER stress, ER synthesized proteins have increased ER retention due to inefficient protein folding in the ER (Mahameed et al., 2020). In our experimental system of CPA-induced ER stress, proinsulin exited the main portion of the ER and formed granule-like structures. The structures did not colocalize with ERGIC and during prolonged stress showed increased localization with the cell periphery (**Figure 2**). Impaired proinsulin processing and proinsulin secretion are hallmarks of T1D and T2D (Kharroubi and Darwish, 2015). The exact nature of these granule-like structures is unknown. However, the correlation with the decreased maturation of PCSK2, the protease that cleaves proinsulin to produce the mature insulin (Bennett et al., 1992), and the decreased lysosomal activity, a mechanism of clearance of insulin granules (Zhou et al., 2020) may implicate these events as part of the mechanism. Additional possibilities could include changes in luminal acidification (Tompkins et al., 2002) and decreased cholesterol in immature secretory granule membranes (Hussain et al., 2018). Furthermore, dysfunction of the zinc transporter, SLC30A8, a known autoantigen in T1D, can also impact on proinsulin to insulin processing (Wijesekara et al., 2010). Interestingly, cholesterol biosynthesis (**Supp. Fig 6**) and the RNA levels of SLC30A8 decreased during chronic ER stress in MIN6 cells, supporting multiple potential contributions to the inhibition of proinsulin processing to insulin.

β-cell rest is described as suppression of insulin release from β-cells, which was originally shown in the context of T1D (Brown and Rother, 2008). Exogenous administration of insulin reduced the overload of endogenous insulin secretion by reducing exogenous insulin requirements suggesting that decreased demand on β-cells can improve insulin secretion and β-cell viability at least transiently. In a similar vein, our findings suggest that the CPA-induced ER exit of proinsulin could decrease β-cell ER load and help cell survival under stress conditions. Molecular insights into the basis of this subcellular relocalization-based β-cell stress mitigation need further investigation. The loss of expression of key β-cell maturation marker genes and impaired insulin secretion have been observed in mouse models of T1D and T2D, as well as in patients with diabetes. It has recently been reported that insulin in the serum of patients with long-term T1D originates from these “dedifferentiated” β-cells (Lam et al., 2019). In line with this, histological evidence provides further support for β-cell dedifferentiation at T1D onset (Seiron et al., 2019). Preclinical and clinical findings from patients with T1D and T2D suggest that a transient recovery of β-cell dysfunction occurs upon glycemia normalization by intensive insulin treatment. It is hypothesized that β-cell rest and/or re-differentiation played a role in the glycemic normalization (Harrison et al., 2012; Wang et al., 2014). These intriguing findings suggest that resting, immature and dedifferentiated states of β-cells could be a part of a unique adaptive mechanism allowing β-cells to camouflage/protect themselves from ongoing stress and potentially escape immune recognition and destruction. Thus, triggering the re-differentiation of dedifferentiated β-cells and restoring their functional capacity may represent a novel, feasible, and important approach for the treatment of diabetes. In that regard, better understanding of the mechanisms of βEAR development can be instrumental for designing novel therapies.

Our studies in MIN6 can be considered as a novel methodology to identify biomarkers for T1D and T2D pathogenesis. Most of the 11 genes identified here (**Sup.Fig. 4A**), associate with T1D and T2D, suggesting a role of ER stress adaptation in the diabetic homeostatic state. For example, G6PC2 is an islet-specific glucose-6-phosphatase catalytic subunit-related protein that catalyzes the hydrolysis of glucose-6-phosphate in both gluconeogenic and glycogenolytic pathways. Genome-wide association studies have identified the association of G6PC2 with both diabetes mellitus and cardiovascular-associated mortality (Bouatia-Naji et al., 2008; Chen et al., 2008). The transcription factor NKX2.2 is a known β-cell identity gene. NKX2.2 is required for the mature β-cell maintenance; dysfunction of NKX2.2 causes downregulation of MAFA and GLUT2, as well as disrupted islet architecture (Doyle and Sussel, 2007).

Remarkably, seven out of the 11 signature genes are associated with either insulin maturation or GSIS. PCSK1/3, as well as PCSK2 are involved in the C-peptide removal from proinsulin, a crucial role in insulin maturation (Davidson et al., 1988). Deficiency of three plasma membrane proteins, GLP1R, ABCC8 and P2RY1 are tightly linked to abnormal GSIS and are associated with diabetes (Fehmann and Habener, 1991; Koufakis et al., 2019; Leon et al., 2005). Two secreted proteins, PPP1R1A and SCGN, showed decreased levels in both ER stress adaptation in MIN6 cells and T1D patients. PPP1R1A is an inhibitor of the protein-phosphatase 1, and is positively correlated with insulin secretion and negatively with HbA1c (Taneera et al., 2015). SCGN is an insulin-interacting calcium sensor protein that stabilizes insulin (Sharma et al., 2019). Due to the secretion property of these two proteins, they have been proposed as biomarkers for diabetes detection (Jiang et al., 2013; Sharma et al., 2019). SYT4, Synaptotagmin 4, localizes in mature insulin granules and is important for GSIS; the overexpression increases GSIS threshold, while the ablation compromises GSIS (Huang et al., 2018). Finally, we implicate a new diabetic candidate gene VAL1L, also known as Vesicle Amine Transport 1-Like, associated with ER stress adaptation in MIN6 cells and T1D. Overall, our data suggest that the use of the genome-wide cellular response in MIN6 cells can lead to the discovery of biomarkers and mechanisms of pathogenesis in diabetes.

Although our current study modeled changes typical of pre-T1D, alternative experimental systems can study T2D pathogenesis (Swisa et al., 2017). For example, lipotoxicity can induce ER stress in tissue cultured β-cells (Cunha et al., 2008), and is a factor associated with β-cell loss in T2D (Cnop et al., 2012). Similar to the studies presented here via the use of a chemical ER stressor, palmitate-induced ER stress in MIN6 cells, results in a similar decrease in MAFA protein levels (**Sup. Fig.7**). Future studies can determine the commonalities between different physiological and pathological stressors and the bona fide ER stress cellular response described here.

Finally, while in this study we take the advantage of a reversible ER stress inducer, it is well-established that ER stress can be induced by pro-inflammatory cytokines, nitric oxide, and reactive oxygen species all of which can cause β-cell damage in T1D (Brozzi and Eizirik, 2016). Within the inflammatory islet microenvironment vicious cycles of cytokine production and unresolved ER stress can lead β-cells to undergo a dormant state. Hence, future experiments by mimicking this islet microenvironment in cultured mouse and human islets would provide an invaluable insight into states of reversibility, regulation and function of βEAR and pave the way for novel approaches preventing the loss of functional β-cell mass in T1D.

## Acknowledgement

We thank Jing Wu for technical assistance during the entire study. This manuscript used data acquired from the Human Pancreas Analysis Program (HPAP-RRID:SCR_016202) Database (https://hpap.pmacs.upenn.edu), a Human Islet Research Network (RRID:SCR_014393) consortium (UC4-DK-112217, U01-DK-123594, UC4-DK-112232, and U01-DK-123716).

## Author contributions

C.-W. C and M.H. provided the conceptual framework for the study. C-W.C. designed the experiments. C.-W.C., B.-J. G., M.R.A, Z.G. and S.B. performed the experiments and collected the data. C.-W.C., H.L., T.L., performed bioinformatics analysis of MIN6 cells and MEFs. C.-W.C., C.E.M. and I.C.G. performed microarray analysis. C.-W.C., L.G. and K.H.K. performed scRNA-seq analysis and assisted with evaluation of the data. C.-W.C., L.H., A.E.S., A.P., B.Z., F.E. and M.H. interpreted the results. M.H. and F.E supervised the project with input from C.-W. C. and B.T. M.H., F.E. and B.T. wrote the paper with input from all authors.

## Conflicts of Interests

The authors declare that they have no competing interests.

## Funding

DK053307 and DK060596 (to MH), DK48280 and DK127747 (to PA). JDRF and Hensley Trust, and UC4DK104155 (to CM).

## Supplementary Figure Legends

**Supp. Figure 1. Insulin levels do not change during CPA treatment and CPA washout.** (A) Western blot analysis of UPR markers in MIN6 cells expressing shRNAs for CON or the eIF4E mRNA and treated with CPA for the indicated times (B) Western blot analysis of insulin by non-reducing PAGE electrophoresis. Arrows indicated either insulin monomer or hexamer.

**Supp. Figure 2. Quality controls and validations of the ribosome foot printing and RNA-seq datasets**. (A-B). Overview of experimental setup. Transcriptomes were generated from poly(A) selected RNA and translatomes from ribosome protected RNA fragments. Reading frame analysis (C) and RNA sequencing reads distribution (D) across the *Ins*2 mRNA. (E) Scatterplots represent the reproducibility of transcriptomes and translatomes in two biological replicates. (F) Gene expressions changes in the transcriptomics dataset from MIN6 cells treated as in A. (G) Relative expression levels for the indicated mRNAs, measured by qRT-PCR. *n*=3 biological replicates. Error bars represent S.E.M. *. P<0.05, **. P<0.01, ***. P<0.005 *two-tailed* paired Students’ *t*-test.

**Supp. Figure 3. Comparative analysis of transcriptomes and translatomes between MIN6 and MEFs in response to ER stress**. (A) Scatterplots of fold changes in MIN6 (*left)*, and MEFS (*right*), during progression from acute to chronic ER stress (CPA18 vs CPA1 and Tg16 vs Tg1, respectively). KEGG pathway analysis of genes with either decreased mRNA abundance (B) or increased mRNA abundance (C) in MIN6 (*upper)* and MEFs (*lower)* panels. (D) Gene ontology analysis for genes in the KEEG pathway “protein processing in ER” in MIN6 and MEFs. Notice, MIN6 cells have 68 genes with increased mRNA abundance (panel C, upper) and 10 genes with increased ribo^ocp^ (panel A, included in the 603 genes).

**Supp. Figure 4.** β-**cell ER stress 334 gene set expression in T1D/T2D datasets**. (A) Relative expression of genes in T1D (*upper*), and T2D (*lowe*r), islet microarray datasets. Genes showing higher decrease in expression (magenta) (log_2_ < 1, T1D or T2D vs CON) or moderate decrease in expression (green) (0.5 < log_2_ ≦ 1, T1D or T2D vs CON) are highlighted. (B) Description of healthy donors and T1D patients who donated islets for the scRNA-seq analysis. (C) violin-plots of genes in β-cells scRNA-seq datasets.

**Supp. Figure 5. MIN6 cells can recover even after prolonged ER stress exposure**. (A) Experimental setup. (B) Western blot analysis for the indicated proteins and treatment of MIN6 cells as shown in A. (C) cell viability during recovery from short (6h), chronic (18h) and prolonged (42h), ER stress exposure.

**Supp. Figure 6. Cholesterol biosynthesis rates in MIN6 cells**. Cholesterol biosynthesis was measure by metabolic labeling of the cells with deuterium and analysis of labelled cholesterol by Gas Chromatography-Tandem Mass Spectrometry. Treatments with CPA are indicated.

**Supp. Figure 7. Palmitic acid-treatment of MIN6 cells induces the UPR gene expression and decreases MAFA protein levels**. Western blot analysis for the indicated proteins in MIN6 cells treated with oleic acid (400 mM), palmitic acid (400 mM) or a 2:1 ration of palmitic acid:oleic acid (400 mM) for 48 h. Analysis of extracts isolated from CPA-treated MIN6 cells is also shown.

## STAR Methods

### Experimental Model and Subject Details

#### Cell Lines, Cell Culture Conditions

Mouse insulinoma MIN6 cells were cultured in DMEM containing 4.5g/L glucose, 10% heat-inactivated FBS, 1mM sodium pyruvate, 0.1% β-mercaptoethanol, 2mM L-glutamine and 100U/mL of Penicillin and Streptomycin. Rat insulinoma INS1E cells were cultured in RPMI 1640 containing 2g/L glucose, 10% FBS, 10mM HEPES, 0.1% β-mercaptoethanol, 2mM L-glutamine and 100U/mL of Penicillin and Streptomycin. Human EndoC-βH3 cells were cultured in DMEM containing 5.6 mM glucose, 2% BSA fraction V, 10mM nicotinamide, 5.5μg/mL transferrin, 6.7ng/mL sodium selenite, 50 μM 2-mercaptoethanol, 100 U/mL Penicillin, 100 U/mL Streptomycin and 10 μg/mL puromycin. Cells were cultured at 37°C, supplied with 5% CO_2_. MIN6 and INS1E cells were subcultured in a 1-to-4 ratio weekly. EndoC-βH3 cells were subcultured in a 1-to-2 ratio biweekly. For shRNA knockdown experiments, lentiviral particles expressing shRNA against the eIF4E mRNA was prepared and propagated in HEK293T cells as described previously (Krokowski et al., 2013) using the second generation pLKO.1, psPAX2 and pMD2.G vectors. After two rounds of lentiviral infection, cells were selected under puromycin (10 μg/mL) for three days.

#### Chemicals, reagents and antibodies

Chemicals used in this study: CPA (200μM, MIN6; 50μM, INS1E; 500μM, EndoC βH3) (Tocris). Cycloheximide (100μg/ml) (Sigma-Aldrich). MG132 (10μM, Sigma-Aldrich) and Bafilomycin A1 (400 nM, APExBIO) were used for the designated times. Palmitic acid (400mM) (Sigma-Aldrich), Oleic acid (400mM) (Sigma-Aldrich). CellTiter-GloR Luminescent Cell Viability Assay (Promega) was used to measure cell viability according to manual. Lysosomal Intracellular Activity Assay Kit (Abcam) was used to measure lysosomal activity according to user manual. Antibodies for Western blot analysis: Rabbit anti-ATF4 (1:1000, Cell Signaling); Rabbit anti-BiP(GRP78) (1:1000, Cell Signaling); Mouse anti-CHOP (1:1000, Cell Signaling); Mouse anti-eIF2α (1:1000, Santa Cruz); Rabbit anti-eIF2α-phospho (Ser51) (1:1000, Abcam); Rabbit anti-GADD34 (1:1000, Santa Cruz); Rabbit anti-PERK (1:1000, Cell Signaling); Rabbit anti-HERPUD1 (1:2000, Cell Signaling); Mouse anti-β-TUBULIN (1:4000, Sigma-Aldrich); Mouse anti-Proinsulin (1:1000, Novus Biologials); Rabbit anti-MAFA (1:2000, Cell Signaling); Rabbit anti-PCSK2 (1:1000, Cell Signaling); Rabbit anti-PDX1 (1:2000, Cell Signaling); Rabbit anti-NKX2.2 (1:1000, Invitrogen); Guinea pig anti-p62(SQSTM1) (1:1000, Progen Biotechnik); Rabbit anti-LC3(ATG8) (1:1000, Novus biologicals); Guinea pig anti-mouse INS serum (1:200) was kindly gifted from Dr Peter Arvan (University of Michigan). Immunofluorescence staining antibodies: Rabbit anti-PDI(P4HB) (1:200, Sigma-Aldrich); Rabbit anti-ERGIC53 (1:200, Sigma-Aldrich); Goat anti-rabbit IgG Alexa-488 (1:200, Invitrogen); Goat anti-mouse IgG Alexa-594 (1:200, Invitrogen); Hoechst33324 (8 μM, Invitrogen).

#### Protein sample preparation for Western blot analysis

For protein extraction, cells were washed twice with ice-cooled 1xPBS before lysis. Ice-cooled lysis buffer (50 mM Tris-HCl pH7.5, 150 mM NaCl, 2mM EDTA, 1% NP-40, 0.1% SDS, 0.5% sodium deoxycholate, supplemented with EDTA-free protease inhibitor (Roche Applied Science) and PhosSTOP phosphatase inhibitor (Roche Applied Science) was added to cells. Cells were scraped off and sonicated on ice. Protein lysates were centrifuged for 5 min in 10000xg at 4°C. Supernatant was collected and quantified by DC Protein Assay Kit (Bio-Rad). The lysate was diluted to 1μg/μL by using lysis buffer. The diluted lysates were mixed with 5x sampling buffer (300mM Tris-HCl pH6.8, 50% glycerol, 10%(v/v) β-mercaptoethanol, 10%(w/v) SDS and 50mg bromophenol blue) or Pierce Lane Marker Non-Reducing Sample Buffer (Thermo Scientific) in a 1-to-4 ratio for reducing or non-reducing Western blot analysis, respectively. For non-reducing analysis, gels were soaked with 25mM DTT for 10 min before electrotransfer to Immobilon-P PVDF membrane (Sigma-Aldrich) (Arunagiri et al., 2019).

#### Measurement of protein synthesis rate

Protein synthesis rates were measured as previously described (Guan et al., 2017). In brief, cells were treated with CPA for the indicated times. At the end of treatments, [35S]Met/Cys (30μCi/mL EXPRE35S Protein Labeling Mix (PerkinElmer)) was added to the cells for an additional 30 min. After labeling, cells were washed and lysed, and the radioactivity incorporated into proteins was determined by liquid scintillation counting. The protein synthesis rate was calculated as the rate of [35S]Met/Cys incorporation to total cellular protein from the same lysate.

#### Measuring *in vitro* guanine nucleotide exchange factor (GEF) activity of eIF2B

EIF2B activity was measured as previously described (Guan et al., 2017; Kimball et al., 1989). In brief, cells were washed and scraped off in homogenization buffer (45 mM HEPES-KOH pH 7.4, 0.375 mM MaOAc, 75 mM EDTA, 95 mM KOAc, 10% glycerol, 1mM DTT, 2.5 mg/mL digitonin, supplemented with EDTA-free protease inhibitor (Roche Applied Science) and PhosSTOP phosphatase inhibitor (Roche Applied Science)). Cell lysates were homogenized and quantified for protein concentration. The EIF2B activity was calculated as the rate of exchange from eIF2α[^3^H]GDP to nonradioactive GDP over the time points.

#### RNA preparation and qRT-PCR

Cells were washed twice with ice-cooled 1xPBS on ice after the designated treatments. Total RNA was extracted using TRIzol LS reagent (Invitrogen). The cDNA was synthesized using SuperScript III First-Strand Synthesis SuperMix (ThermoFisher). The relative quantity of specific mRNAs was measured by using VeriQuest SYBR Green qPCR Master Mix (ThermoFisher) with StepOnePlus Real-Time PCR System (Applied Biosystem). Primers used in this study are listed in **Supp.Table 5**.

#### Immunofluorescence staining

MIN6 and INS1E cells, were plated on glass microscope cover slips (ThermoFisher) in 6cm culture dishes and were allowed to grow for 48 hours. For Human EndoC- βH3 cells, cells were plated on coated (matrigel (1:500), Sigma-Aldrich and fibronectin (1μg/cm2), Sigma-Aldrich) glass microscope cover slips (ThermoFisher) in 6cm culture dishes and allowed to grow for 1 week. After the designated treatments, cells were washed twice with ice-cooled 1xPBS on ice. Cells were fixed with 4% paraformaldehyde for 10 min. Fixed cells were washed twice with ice-cooled 1xPBS and incubated in PBST (1xPBS + 0.02% Triton X-100) for 15 min, PBST with 10% FBS for 30 mins, and PBST with 10% FBS and primary antibodies at 4°C for 16h. After washing with ice-cooled PBST twice, cells were incubated in PBST with 10% FBS and secondary antibodies for 2 h in dark. This was followed by washing with ice-cooled PBST twice and nuclei staining with Hoechst 33342 for 5 min in dark. After washing with ice-cooled PBST twice, cells were mounted in Fluoromount-G (Electronic Microscope Sciences) and sealed with clear nail polish on microscope slides. The images were captured using a Leica SP8 confocal microscope. Imaging areas were randomly selected in a single-blind manner by a microscope specialist. Four imaging areas were fetched in each condition, while a representative image was shown.

#### Deep sequencing library generation

The general procedures for generation of mRNA-seq and Ribo-seq libraries were previously reported by us and others (Chen and Tanaka, 2018; Ingolia et al., 2009). In brief, cells were washed twice by ice-cooled PBS with 100μg/ml cycloheximide (Sigma-Aldrich) before harvest. Lysis buffer (10 mM Tris, pH 7.4, 100 mM NaCl, 140mM KCl, 10mM MgCl_2_, 1 mM DTT, 1% (v/v) Triton X-100, 500 U/mL RNasin and 100 μg/mL cycloheximide) was added and cells were scraped fast on ice. The lysate was centrifuged for 5 min in 500xg at 4°C. The supernatant was collected and centrifuged for 10 minutes in 10000xg at 4°C. The supernatant was collected for mRNA and monosome enrichments. For mRNA enrichment, total RNA was extracted from lysates with TRIzol LS reagent (Invitrogen). The RNA with poly adenosine tract was purified by a Magnetic mRNA Isolation Kit (New England Biolabs) according to the user manual. The RNA with poly adenosine tract was fragmented with an alkaline buffer (2 mM EDTA, 10 mM Na_2_CO_3_, 90 mM NaHCO_3_, pH ≈ 9.3) treatment for 40 min at 95°C. For monosome enrichment, lysates were treated with RNase If (50U/100μg, New England Biolabs) for 40 mins at 25°C. Ice-cooled open top 10-50% sucrose gradient centrifuge tubes (14 x 89mm, SETON) were generated by Gradient master 108 (Biocomp). The digested lysate was loaded on top of the gradients and were centrifuged in a SW41Ti rotor (Beckman) for 2 h in 40000 rpm at 4°C in an L-70 ultracentrifuge (Beckman). The monosome enriched fraction was collected by a PGFip Psiton Gradient Fractionator (Biocomp). Total RNA was purified and extracted by TRIzol LS reagent. RNA length between 20 to 40 bases was size selected by 15% TBU gel (Invitrogen). The 3’ ends of the selected RNA fragments were dephosphorylated by T4 PNK (New England Biolabs). Ribosomal RNA was removed from the RNA isolated from monosome enrichment, by NEBnext rRNA depletion kit (New England Biolabs). RNA was ligated with Linker-A (New England Biolabs) by T4 RNA ligase II KQ (New England Biolabs). Ligated RNA fragments were size selected by 10% TBU gels (Invitrogen). Selected RNA was reverse transcribed (RT) with in-house-designed DNA oligos (**Supp.Table 5**) by ReverTra Ace (Toyobo) according to the user manual. The RT product was circularized with CircLigase II ssDNA Ligase (Lucigen). The libraries were amplified by Phusion High-Fidelity DNA Polymerase (New England Biolabs) for 10 polymerase-chain-reaction (PCR) cycles with in-house-designed indexed PCR oligos (**Supp. Table 5**). PCR products between 140 to 170 bps were collected by 4-20% TBE gel (Invitrogen) for deep sequencing. The libraries were quantified and sequenced by Novogene Corporation in Illumina HiSeq platform. Three independent biological replicates were performed for bioinformatics analysis.

#### Islet microdissection and microarray analysis

The microarray data preparation was described previously (Holm et al., 2018; Trembath et al., 2020). In brief, frozen tissue was obtained from the Network for Pancreatic Organ donors with Diabetes (nPOD) (**Supp. Table 4**) (Pugliese et al., 2014). Optimal cutting temperature slides of pancreatic tissue were used for laser-capture microscopy that was conducted on an Arcturus Pixcell II laser capture microdissection system (Arcturus Bioscience). All islets visible in two to five sections from each sample were pooled, and RNA was extracted using the Arcturus PicoPure RNA Isolation Kit (Applied Biosystems). RNA quantity and quality were determined using a Bioanalyzer 2100 (Agilent Technologies). RNA samples were subjected to gene expression analysis using Affymetrix expression arrays (ThermoFisher). Fluorescence intensity was used to refer to gene expression.

#### Single-cell RNA sequencing analysis

The scRNA-seq raw data from T1D and healthy donors datasets were obtained from HPAP (https://hpap.pmacs.upenn.edu/) (Kaestner et al., 2019). The raw data were pooled and normalization using SCtransform (Hafemeister and Satija, 2019) in the Seurat package in R. No additional batch-effect removal step was applied. Cell types were determined by Seurat clustering results according to the expression of pancreatic marker genes (Muraro et al., 2016). The cell profiling was done by a machine learning approach via the Rtsne package in R according to the expression of the UPR signature (GO:0030968) and 6 β-cell identity genes, *INS*, *PAX6*, *NKX2.2*, *NKX6.1*, *MAFA* and *PDX1*. Patients’ information used in this study was listed in Sup. Fig. 4B.

#### Cholesterol synthesis measured by GC-MS

Cholesterol synthesis was determined following deuterium labeling incorporation from deuterium-enriched media. In brief, MIN6 cells were incubated in the medium containing 10% deuterated water (^2^H_2_O, molar percent enrichment) with CPA for 18h (CPA18); while without CPA served as control. For the recovery sample, after 18h of CPA treatment, the cells were washed twice with 1xPBS and refreshed with the medium containing 10% deuterated water for 24h incubation. After incubation, the supernatant was collected for determination of deuterium enrichment in the medium. After 1xPBS wash twice, cells were trypsinized and pelleted by centrifugation at 4°C at 650xg for 5 min. Cell pellets were washed with 1xPBS and pelleted by centrifugation at 4°C at 650xg for 5 min. Supernatants were discarded and pre-cooled methanol was added into the tubes containing cell pellets. After mixing by inversion for 5 times, cells were pelleted by centrifugation at 4°C at 650xg for 5 min. The supernatant was discarded and pellets were stored at -80°C until extraction.

The ^2^H-labeling of body water was determined by the exchange with acetone, as previously described by (Katanik et al., 2003) and further modified. In brief, 5 µL of cellular medium or standard were reacted for 4 h at room temperature with 5 µL of 10 N KOH and 5 µL of acetone. Headspace samples were injected into GC/MS in split mode (1:40). Acetone m/z 58 and 59 were monitored. Isotopic enrichment was determined as ratio of m/z 59/(58+59) and corrected using a standard curve.

For cholesterol extraction, cell pellets in the tubes were homogenized frozen in 600 µl of Folch solution (chloroform:methanol, 2:1, v/v) on dry ice. After addition of 0.4 volumes of ice-cold water, cells were homogenized again and let incubate on ice for 30 min. Homogenates were centrifuged at 4°C at 14,000 rpm for 10 min. Upper methanol/water layer was discarded. Internal standard (heptadecanoic acid) was added to the bottom chloroform layer and was evaporated to dryness. Cholesterol was then converted to TMS (trimethylsilyl) derivative by reacting with bis(trimethylsilyl) trifluoroacetamide with 10% trimethylchlorosilane (Regisil) at 60°C for 20 min. Resulting TMS derivatives were run in GC-MS.

GC-MS conditions used for the analysis were carried out on an Agilent 5973 mass spectrometer equipped with 6890 Gas Chromatograph. A DB17-MS capillary column (30 m × 0.25 mm × 0.25 μm) was used in all assays with a helium flow of 1 mL/min. Samples were analyzed in Selected Ion Monitoring (SIM) mode using electron impact ionization (EI). Ion dwell time was set to 10 msecs. Cholesterol m/z 368-372 were monitored to detect isotopic label incorporation.

The absolute cholesterol biosynthesis was computed as follows. Cholesterol GC-MS data were corrected for natural abundance and resulting enrichments were divided into precursor isotopic enrichment of media deuterium (10%) to follow precursor-product relationship. This resulted in fractional synthesis rate. Absolute cholesterol synthesis rate was determined by multiplying fractional rate by the cholesterol content which was determined using internal standards.

#### Biostatistics and bioinformatics

Mouse RNA reference (GRCm38.p6.rna) was used for sequencing reads alignment. The alignment was done by Bowtie2 (Langmead and Salzberg, 2012) in an in-house-established Galaxy platform. In brief, raw sequencing reads were filtered by quality, Q30 > 90%. The length of reads between 15 and 40 were selected. The selected reads were aligned to the RNA reference with parameters of equal or higher than 95% identity and less than one mismatch was allowed. Genes with at least one read aligned and identified in all three biological replicates were collected for further analysis. Transcript count was calculated as reads per kilo-base per million reads of total aligned reads (RPKM). Ribosome occupancy (Ribo^ocp^) of a transcript was calculated as the mean of transcript count in Ribo-seq divided by the mean of transcript count in mRNA-seq. Gene functional annotations were done in KEGG: Kyoto Encyclopedia of Genes and Genomes (https://www.genome.jp/kegg/) and Gene Ontology Resource (http://geneontology.org/). Data analysis, statistics and presentations were done by R (https://www.r-project.org/) and Excel (Microsoft). Bar charts were done by Excel. Heatmaps were done by Heatmap2 package in R. Scatterplots, volcano plots and violin plots were done by the Ggplot2 package in R.

The statistics of the heatmaps in Figure 3 was done by Excel. For Figure 3C, the changes of transcript expression in [log2] values of CPA1 over control (acute), CPA18 over CPA1 (chronic) and CPA18 over control (overall) were calculated. The cutoffs to define whether the transcript was upregulated or downregulated were set as 0.32 or -0.32, respectively. Transcripts that showed upregulation in both acute and chronic phases were grouped into G1. Transcripts that showed upregulation in the acute phase but downregulation in the chronic phase were grouped into G2. Transcripts that showed no change in the acute phase but upregulation in the chronic phase but overall upregulation were grouped into G3. Transcripts that showed downregulation in both, acute and chronic phases were grouped into G4. Transcripts that showed downregulation in the acute phase but upregulation in the chronic phase were grouped into G5. Transcripts that showed no change in the acute phase but downregulation in the chronic phase but overall downregulation were grouped into G6. For Figure 3D, transcripts were grouped separately according to the Ribo^ocp^ values. Transcripts that showed increased Ribo^ocp^ in both acute and chronic phases were grouped into G7. Transcripts that showed increased Ribo^ocp^ in the acute but decreased Ribo^ocp^ in the chronic phase were grouped into G8. Transcripts that showed no change of Ribo^ocp^ in the acute phase and an increase in the chronic phase, leading to overall increased Ribo^ocp^ were grouped into G9. Transcripts that showed no change of Ribo^ocp^ in the acute phase and a decrease in the chronic phase, leading to overall to decreased Ribo^ocp^ were grouped into G10. Transcripts that showed decreased Ribo^ocp^ in the acute, but increased Ribo^ocp^ i in the chronic phase were grouped into G11. Transcripts that showed decreased Ribo^ocp^ in both acute and chronic phases were grouped into G12. The functional enrichments of the 12 groups were analyzed by. the KEGG pathway via the DAVID bioinformatics database. The number of genes identified in a single biological pathway was denoted.

For the transcriptome and translatome analysis the gene expression levels represent the means of RPKM of three biological replicates. The values of differential expression were determined as log2(expression in RPKM in condition 2 – expression in RPKM in condition 1) (L2DE). The cuff-off value for data filtering and statistics was set as log2=0.6 or -0.6, which represented at least 50% difference in RPKMs between two conditions. The definition of expression regulation was set as described previously (Guan et al. 2017). In brief, up-regulation in both mRNA abundance and ribo_ocp_, -0.6≦(x-y)<0.6, x≧0.6, y≧0.6; up-regulation in ribo_ocp_ only, (x-y)<-0.6, y>0.6; down-regulation in both mRNA abundance and ribo_ocp_, -0.6≦(x-y)<0.6, x≦-0.6, y≦ -0.6; down-regulation in ribo_ocp_ only, (x-y)>0.6, y<-0.6, where x represents the L2DE in transcriptomic data and y represents the L2DE in ribosome footprints.

## Notes

### Competing Interest Statement

The authors have declared no competing interest.

