## Supplemental Figures for "Adaptation to chronic ER stress enforces pancreatic β-cell plasticity"

Supplementary Figure 1. Related to Figure 1.

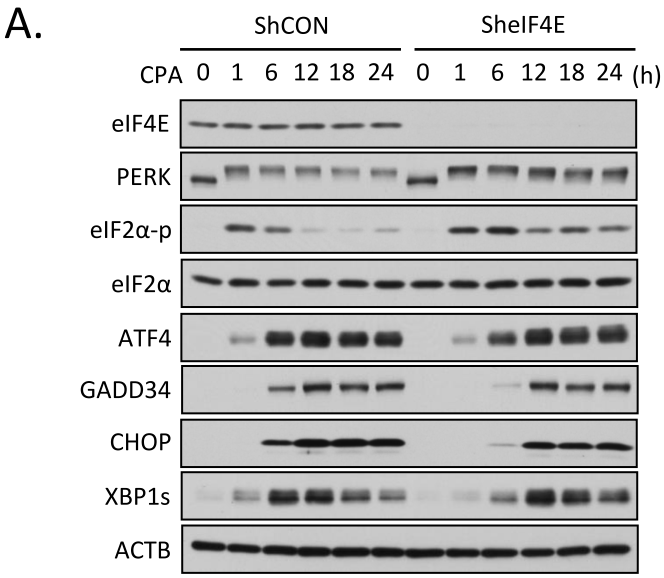


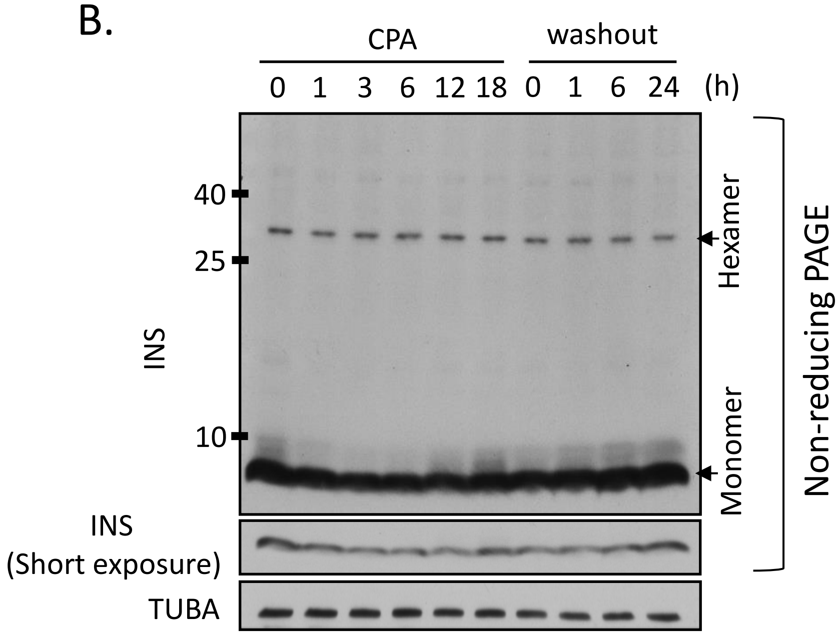

Supplementary Figure 2. Related to Figure 3.

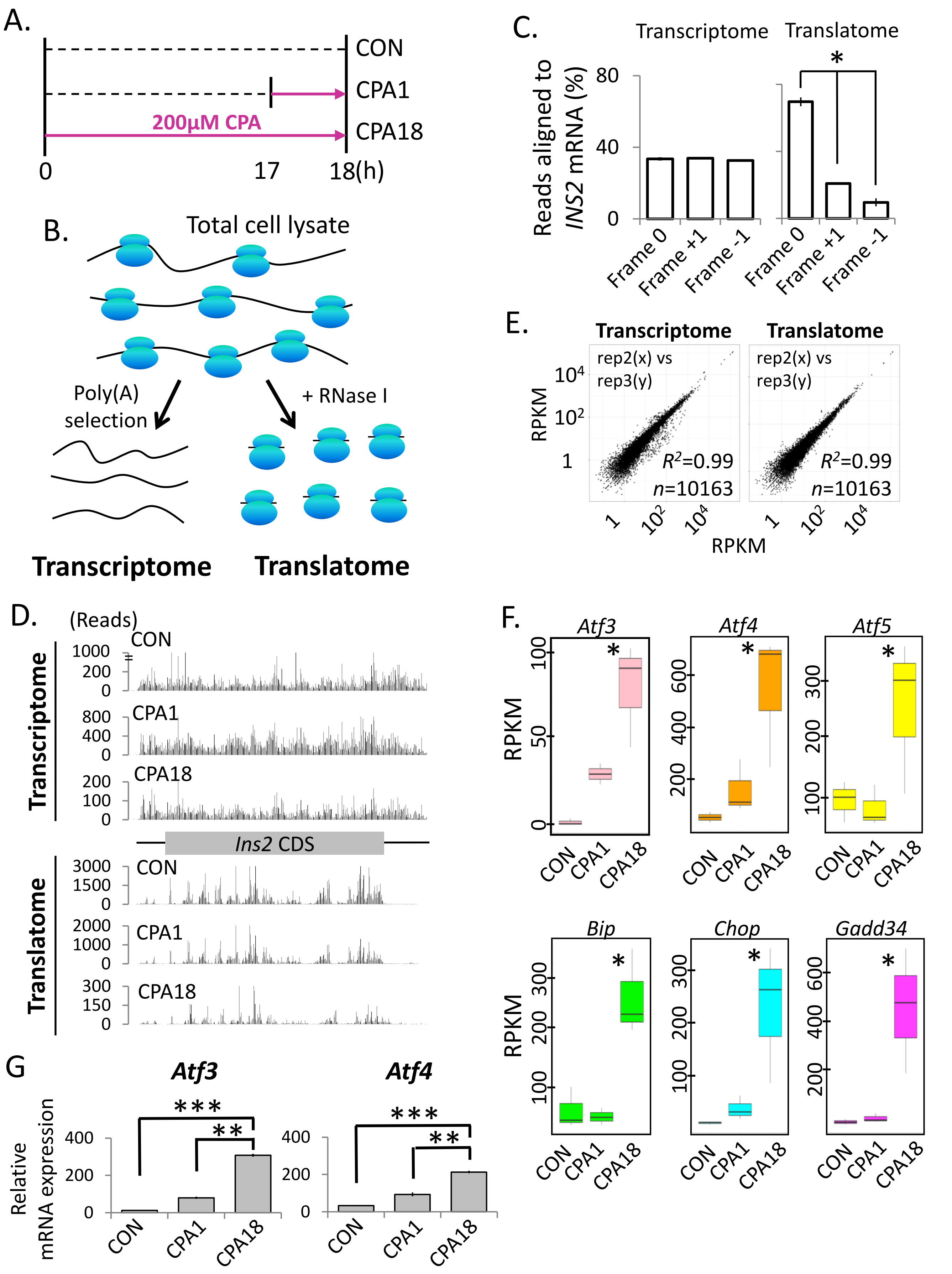

Supplementary Figure 3. Related to Figure 4.

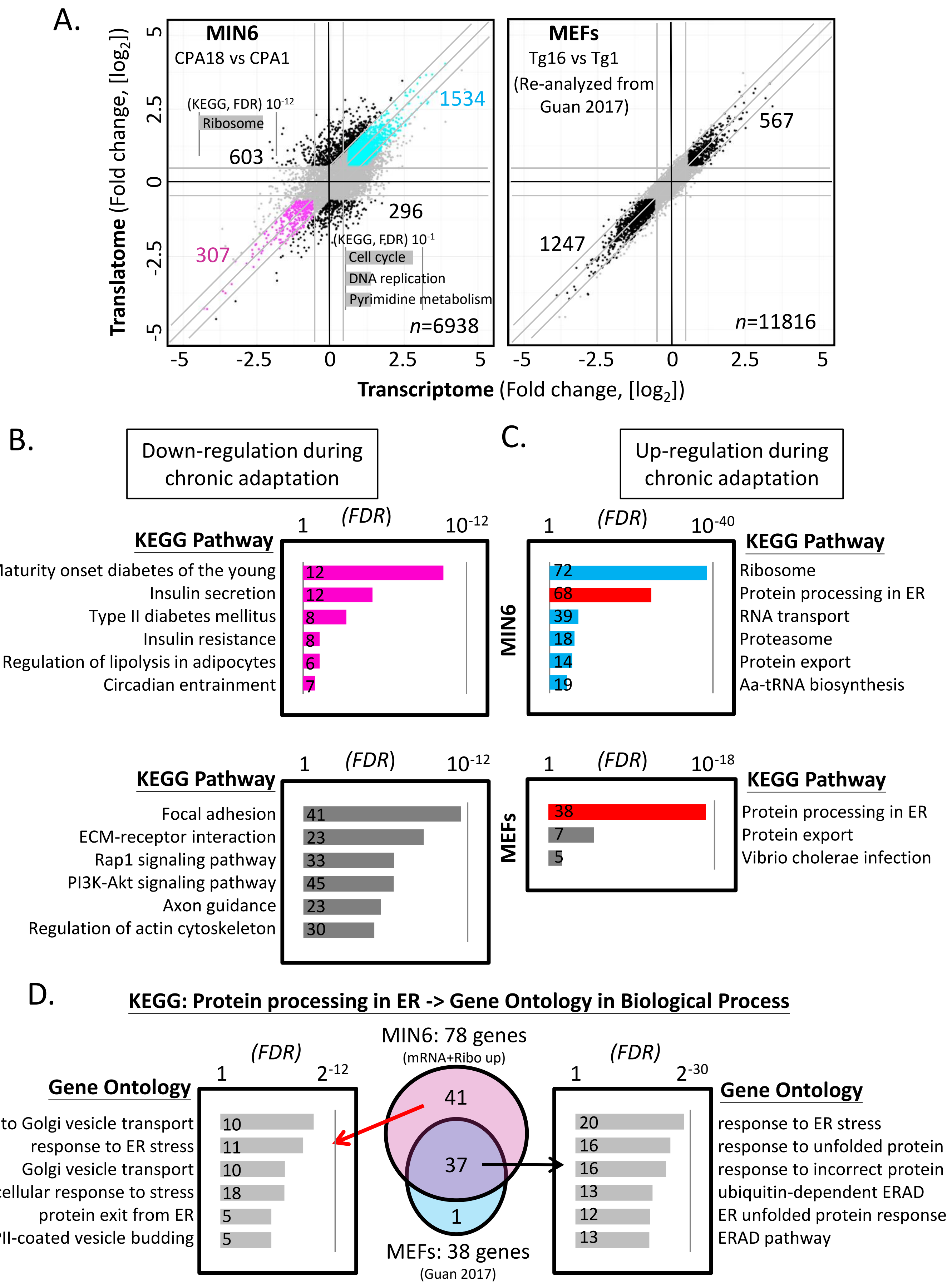

### Supplementary Figure 4. Related to Figure 6.

A.

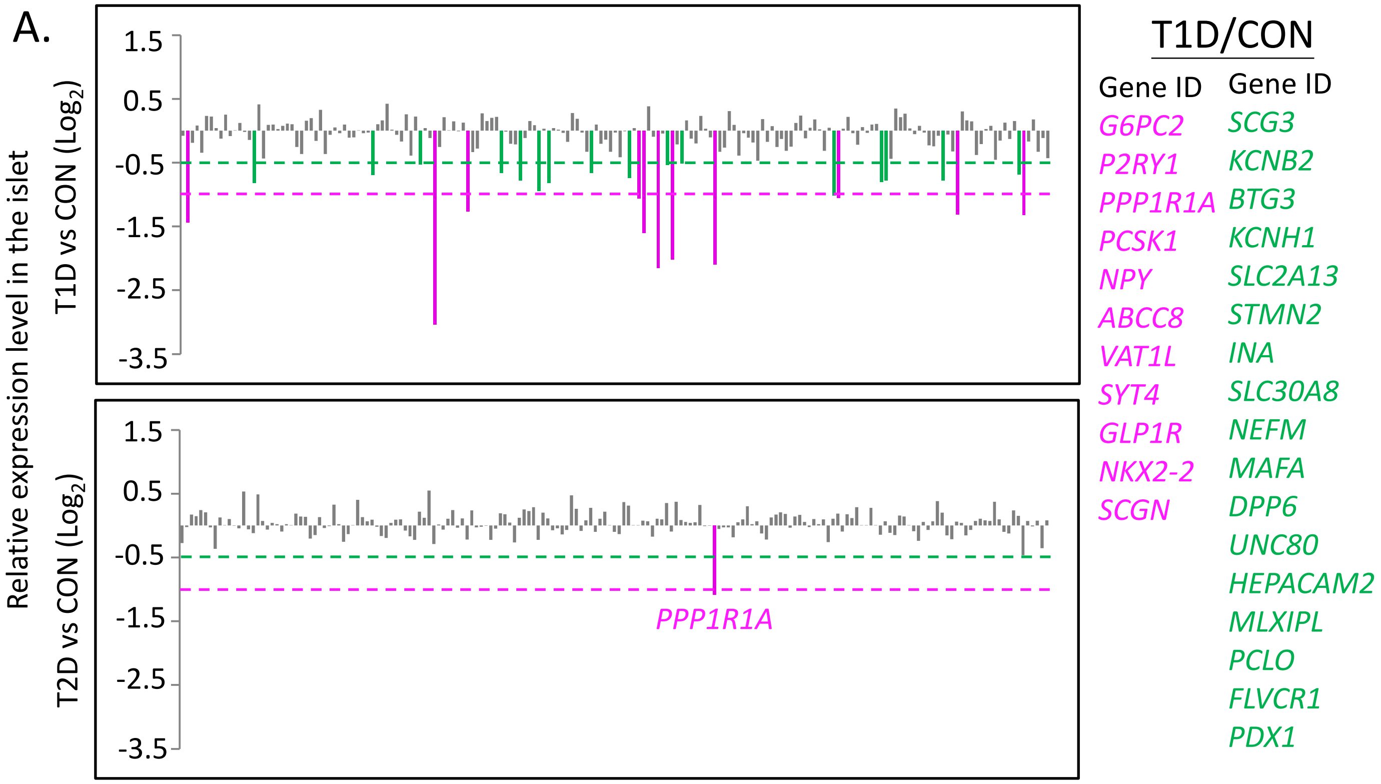


B.

| PancDB No. | Age | Sex | T1D duration | α cell (%) | β cell (%) | δ cell (%) | Pp cell (%) | Acinar cell (%) | Duct cell (%) | Others (%) |
|---|---|---|---|---|---|---|---|---|---|---|
| HPAP019 | 24 | M | ND | 874 (21.51) | 1278 (31.45) | 26 (0.64) | 822 (20.23) | 613 (15.08) | 249 (6.13) | 202 (4.97) |
| HPAP022 | 37 | F | ND | 511 (13.74) | 1069 (28.75) | 70 (1.88) | 73 (1.96) | 380 (10.22) | 1097 (29.51) | 518 (13.93) |
| HPAP024 | 18 | M | ND | 126 (6.78) | 1106 (59.59) | 278 (14.98) | 50 (2.69) | 165 (8.89) | 24 (1.29) | 107 (5.77) |
| HPAP026 | 24 | M | ND | 200 (35.53) | 94 (16.7) | 60 (10.66) | 37 (6.57) | 66 (11.72) | 49 (8.7) | 57 (10.12) |
| HPAP029 | 23 | M | ND | 440 (19.51) | 423 (18.76) | 352 (15.61) | 75 (3.33) | 192 (8.51) | 406 (18) | 367 (16.27) |
| HPAP020 | 14 | M | 5 days | 4319 (49.98) | 1055 (12.21) | 116 (1.34) | 62 (0.72) | 2229 (25.8) | 629 (7.28) | 231 (2.67) |
| HPAP021 | 13 | F | 7 years | 1526 (32.18) | 812 (17.12) | 44 (0.93) | 0 (0) | 1780 (37.54) | 509 (10.73) | 71 (1.5) |
| HPAP023 | 17 | F | 7 years | 300 (34.92) | 86 (10.01) | 10 (1.16) | 1 (0.12) | 215 (25.03) | 171 (19.91) | 76 (8.85) |
| HPAP028 | 4 | M | 3 years | 861 (20.59) | 1407 (33.64) | 394 (9.42) | 31 (0.74) | 365 (8.73) | 735 (17.58) | 389 (9.3) |
| HPAP032 | 10 | F | 3 years | 538 (22.63) | 942 (39.67) | 468 (19.69) | 0 (0) | 217 (9.13) | 47 (1.98) | 165 (6.94) |
| HPAP055 | 24 | M | 7 years | 58 (3.25) | 77 (4.32) | 793 (44.5) | 2 (0.11) | 77 (4.32) | 569 (31.93) | 206 (11.56) |

C.

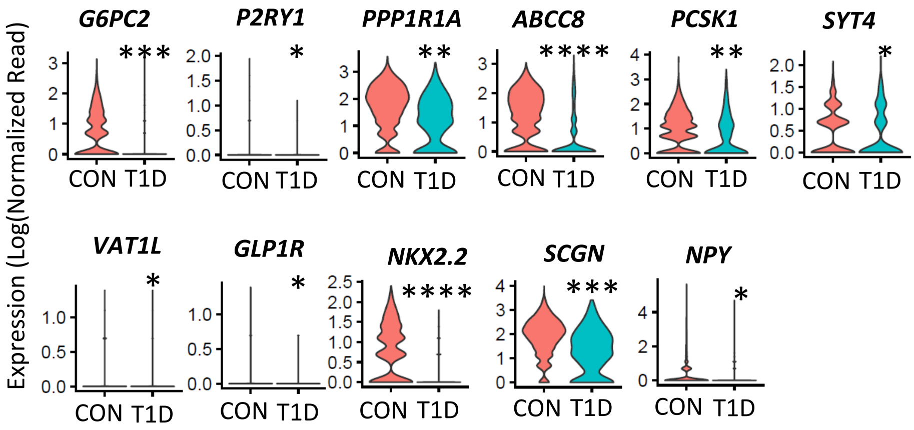

Supplementary Figure 5. Related to Figure 7.

A.

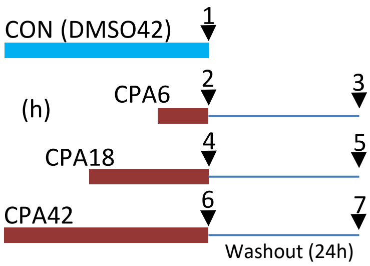


B.

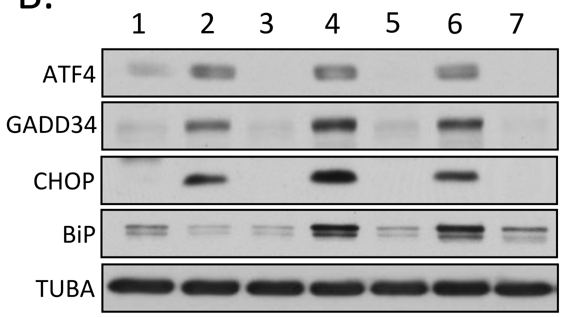


C.

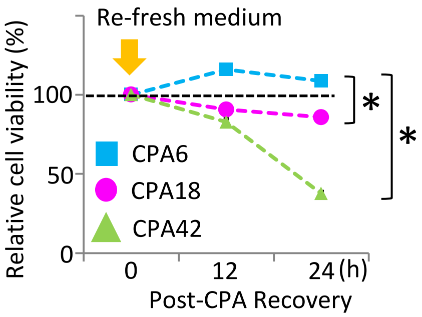

Supplementary Figure 6.

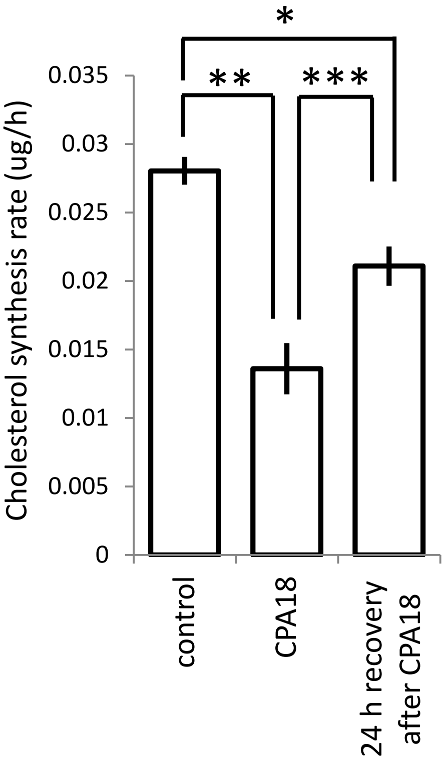

### Supplementary Figure 7.

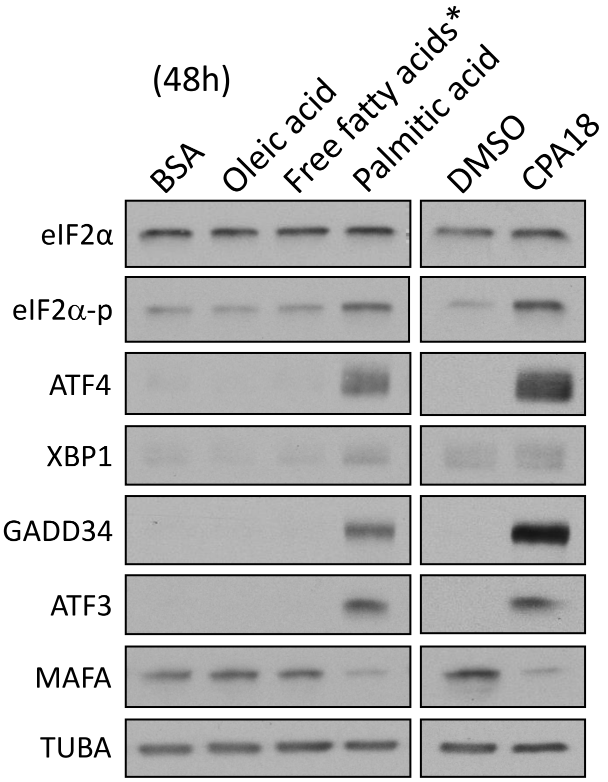


*. Palmitic acid : Oleic acid = 2:1
