## Supplemental Table 2 for "Adaptation to chronic ER stress enforces pancreatic β-cell plasticity"

Supplementary table 2. UPR signature used in this study, related to Figure 3 and 7

| UPR signature (GO:0030968) | MiN6 0vs1 mRNA | MIN6 0vs1 RP | MIN6 0v18 mRNA | MIN6 0v18 RP |
| --- | --- | --- | --- | --- |
| DERL3 | -1.1711968 | 0.78880821 | 2.72968751 | 2.85519369 |
| EIF2A | -1.1641029 | -0.5148726 | -1.4850187 | -0.2959738 |
| TMEM33 | -0.9981014 | -0.1176809 | 0.45347835 | 0.84060865 |
| YOD1 | -0.9565209 | -0.0682751 | -2.0264869 | 0.52326262 |
| CLGN | -0.8725694 | -1.203986 | -1.6530244 | -0.5167689 |
| UFL1 | -0.7766677 | -0.1708048 | 0.43760846 | 0.38249751 |
| DNAJC3 | -0.6909561 | -0.335904 | 0.52751209 | 0.5387983 |
| ATF6 | -0.5037449 | 0.29052198 | 1.22746439 | 2.20913538 |
| EP300 | -0.4814621 | -0.1002759 | -1.9000179 | -0.5810186 |
| ERMP1 | -0.4751891 | -0.5255814 | -0.3262332 | -0.1154211 |
| NFE2L2 | -0.4648596 | 0.4121124 | -1.4176399 | 0.43455956 |
| SERP1 | -0.4031267 | 0.07263977 | 1.2668646 | 2.02267735 |
| VAPB | -0.3483267 | -0.0588688 | 0.60272498 | 0.59342408 |
| HSPA5/BIP | -0.345375 | -0.0345295 | 1.64335475 | 1.94788041 |
| CANX | -0.3404167 | -0.9001733 | -0.4153801 | -0.3566493 |
| DERL1 | -0.2671849 | 0.05129573 | 1.60045491 | 1.51300596 |
| TMED2 | -0.2632033 | -0.3200026 | 0.90897519 | 0.68598065 |
| FICD | -0.2428241 | -0.4556141 | 1.10440685 | 1.03189937 |
| PIK3R1 | -0.2357065 | 0.55479832 | -1.5061506 | 0.31392153 |
| SELENOS | -0.2084829 | 0.11992793 | 1.83990683 | 1.87151794 |
| STUB1 | -0.2065652 | 0.14055631 | 0.60893229 | 0.7312939 |
| WFS1 | -0.1537298 | -0.3715017 | 2.2480869 | 1.83323113 |
| CALR | -0.1518659 | -0.1522317 | 1.59598491 | 1.36098012 |
| AMFR | -0.1291368 | -0.7990096 | 0.74281565 | -0.0567427 |
| ABCA7 | -0.1235075 | -0.1144633 | -0.7011274 | -1.2025298 |
| HERPUD2 | -0.1187214 | -0.9208519 | 0.941012 | 1.03757242 |
| TBL2 | -0.098644 | 0.70128044 | 1.3230328 | 1.27794189 |
| ATF6B | -0.0888714 | -0.2964359 | 1.0169563 | 0.71732419 |
| COPS5 | -0.077204 | -0.1635588 | 0.49978149 | -0.2581285 |
| PDIA6 | -0.0759139 | -0.3061534 | 1.31102581 | 0.86119686 |
| PTPN1 | -0.0707095 | 0.72934935 | 0.70708626 | 0.55820057 |
| ERN1 | -0.036015 | 0.14051458 | 0.7192314 | 1.06255336 |
| CDK5RAP3 | -0.0336151 | 0.20422278 | 1.1706729 | 1.12980092 |
| CREB3 | 0.06585647 | 0.44587831 | 1.74717038 | 1.43498107 |
| DDRGK1 | 0.06585649 | -0.4595818 | 1.66219902 | 0.83708065 |
| EIF2AK3 | 0.12240025 | 0.18818804 | 1.25996875 | 1.38790476 |
| BAX | 0.12380222 | -0.3665485 | 0.68918012 | 0.03170065 |
| XBP1 | 0.31547133 | 1.29785091 | 1.61222897 | 1.80041991 |
| CREB3L2 | 0.32244238 | 0.39848562 | 0.77072872 | 0.53252194 |
| DERL2 | 0.33086493 | 0.1142674 | 1.51855047 | 1.06297276 |
| RNF121 | 0.43405764 | -0.2771743 | 1.57140718 | 0.99564899 |
| STC2 | 0.56596385 | 0.18869241 | 2.57131372 | 2.31789345 |
| BFAR | 0.77335024 | 0.35553346 | 0.77678381 | 2.37890369 |
| PPP1R15A/GADD34 | 0.78225179 | 2.11550941 | 4.56860471 | 4.77938205 |
| HERPUD1 | 1.03222154 | 1.51750776 | 3.07004808 | 2.87679617 |
| ATF4 | 1.43160717 | 2.5666377 | 3.154003 | 3.64067928 |
| PTPN2 | 1.48537646 | 0.83350412 | 0.26319829 | 0.68693683 |
| DDIT3/CHOP | 2.0050082 | 3.75650701 | 4.65585626 | 5.93210412 |
| AGR2 | #N/A | #N/A | #N/A | #N/A |
| ATF3 | #N/A | #N/A | #N/A | #N/A |
| BAK1 | #N/A | #N/A | #N/A | #N/A |
| BHLHA15 | #N/A | #N/A | #N/A | #N/A |
| BOK | #N/A | #N/A | #N/A | #N/A |
| CALR3 | #N/A | #N/A | #N/A | #N/A |
| CALR4 | #N/A | #N/A | #N/A | #N/A |
| CASP12 | #N/A | #N/A | #N/A | #N/A |
| CCND1 | #N/A | #N/A | #N/A | #N/A |
| CREB3L1 | #N/A | #N/A | #N/A | #N/A |
| CREB3L3 | #N/A | #N/A | #N/A | #N/A |
| CREB3L4 | #N/A | #N/A | #N/A | #N/A |
| CREBRF | #N/A | #N/A | #N/A | #N/A |
| EIF2AK2 | #N/A | #N/A | #N/A | #N/A |
| EIF2AK4 | #N/A | #N/A | #N/A | #N/A |
| ERN2 | #N/A | #N/A | #N/A | #N/A |
| ERO1A | #N/A | #N/A | #N/A | #N/A |
| IFNG | #N/A | #N/A | #N/A | #N/A |
| NCK1 | #N/A | #N/A | #N/A | #N/A |
| NCK2 | #N/A | #N/A | #N/A | #N/A |
| PARP16 | #N/A | #N/A | #N/A | #N/A |
| SERP2 | #N/A | #N/A | #N/A | #N/A |
| TMTC4 | #N/A | #N/A | #N/A | #N/A |
