## Supplemental Table 3 for "Adaptation to chronic ER stress enforces pancreatic β-cell plasticity"

Supplementary table 3. BEAR gene identified in this study, related for Figure 7

| BEAR-geneID | MIN6-18v0-mRNA | MIN6-18v0-RP | scRNA-T1D/ctr | scRNA-p-value | scRNA-FDR | microarray-T1D/con | microarray-p-value | microarray-FDR |
| --- | --- | --- | --- | --- | --- | --- | --- | --- |
| NKX2.2 | -0.87243615 | -1.145470139 | -2.120457233 | 1.23E-127 | 9.594E-126 | -1.061708604 | 1.38856E-06 | 7.90411E-06 |
| G6PC2 | -1.904234843 | -1.841019286 | -1.52653142 | 1.67E-80 | 1.86086E-79 | -3.036567307 | 7.43075E-15 | 5.49876E-13 |
| GLP1R | -2.575760216 | -2.538169234 | -1.491011723 | 0.00000595 | 8.14211E-06 | -1.262874975 | 1.60773E-07 | 1.18972E-06 |
| ABCC8 | -1.125247193 | -1.727530221 | -1.437592752 | 4.03E-109 | 1.0478E-107 | -1.434011529 | 9.81203E-12 | 1.81523E-10 |
| P2RY1 | -1.572860442 | -1.445299951 | -1.352764123 | 0.000161515 | 0.000203196 | -2.151281913 | 3.94074E-11 | 5.8323E-10 |
| VAT1L | -0.767422984 | -0.639869983 | -1.230140388 | 3.36E-09 | 5.57617E-09 | -1.324044464 | 1.52432E-08 | 1.41E-07 |
| PCSK1 | -1.730561796 | -1.65409475 | D.Q. | D.Q. | D.Q. | -2.016532978 | 2.85526E-12 | 1.05645E-10 |
| SCGN | -2.092499679 | -2.079033897 | D.Q. | D.Q. | D.Q. | -1.052701639 | 9.66985E-08 | 7.95077E-07 |
| NPY | -1.717892678 | -1.192885397 | D.Q. | D.Q. | D.Q. | -1.603412369 | 0.00200979 | 0.006466281 |
| PPP1R1A | -1.013125316 | -0.9918662 | D.Q. | D.Q. | D.Q. | -2.096515164 | 6.27685E-11 | 7.74145E-10 |
| SYT4 | 0.814462928 | 1.880823074 | D.Q. | D.Q. | D.Q. | -1.313183672 | 9.18722E-12 | 1.81523E-10 |
| KCNB2 | 0.742526125 | 0.602387408 | -1.331652819 | 0.00000777 | 1.04493E-05 | D.Q. | D.Q. | D.Q. |
| SLC2A13 | -1.184035421 | 0.905851349 | -1.211842859 | 1.27E-34 | 5.21368E-34 | D.Q. | D.Q. | D.Q. |
| DPP6 | -1.326756219 | -1.172176726 | -1.214021874 | 2.82E-27 | 8.46E-27 | D.Q. | D.Q. | D.Q. |
| HEPACAM2 | -2.895426512 | -2.659375524 | -2.040030001 | 3.78E-81 | 4.914E-80 | D.Q. | D.Q. | D.Q. |
| MLXIPL | -0.466300298 | -1.548666409 | -1.538774485 | 3.5E-73 | 3.4125E-72 | D.Q. | D.Q. | D.Q. |
| PCLO | -1.515475043 | -0.984282893 | -1.287600947 | 1.11E-32 | 4.12286E-32 | D.Q. | D.Q. | D.Q. |
| RCAN2 | -0.84749366 | -0.815640336 | -1.079541746 | 9.97E-31 | 3.38113E-30 | D.Q. | D.Q. | D.Q. |
| SLC37A1 | 0.03129344 | 0.765893866 | -1.059293688 | 8.88E-10 | 1.5392E-09 | D.Q. | D.Q. | D.Q. |
| CACNA1H | 0.493206012 | -0.956102882 | -2.257412507 | 6.15E-25 | 1.65414E-24 | D.Q. | D.Q. | D.Q. |
| MTMR7 | -0.628175689 | -0.6203879 | -1.30313883 | 3.16E-08 | 4.9296E-08 | D.Q. | D.Q. | D.Q. |
| TMEM151A | 0.076094621 | 0.859391134 | -1.073590428 | 7.55E-10 | 1.33841E-09 | D.Q. | D.Q. | D.Q. |
| CTNNA2 | -1.130205428 | 0.689924342 | -1.104943462 | 0.00000233 | 3.24536E-06 | D.Q. | D.Q. | D.Q. |
| MRPL53 | 0.734858828 | 0.997216521 | -1.883090853 | 2.79E-29 | 9.0675E-29 | D.Q. | D.Q. | D.Q. |
| PPP1R3E | 0.063680991 | 0.729082397 | -2.549331375 | 6.58E-102 | 1.2831E-100 | D.Q. | D.Q. | D.Q. |
| MAST1 | -0.375440862 | -1.353014517 | -1.722237447 | 0.000000219 | 3.22302E-07 | D.Q. | D.Q. | D.Q. |
| SYNE4 | 1.215436856 | 0.848160663 | -2.026621976 | 1.08E-65 | 8.424E-65 | D.Q. | D.Q. | D.Q. |
| RGS7BP | -1.582603443 | 0.873563268 | -1.563064031 | 0.133432578 | 0.136943962 | D.Q. | D.Q. | D.Q. |
| GPR27 | 0.0673469 | 1.296701637 | -1.050487059 | 1.62E-17 | 3.61029E-17 | D.Q. | D.Q. | D.Q. |
| PPP2R2C | -0.548759661 | 0.61583188 | -2.127701346 | 1.13E-26 | 3.26444E-26 | D.Q. | D.Q. | D.Q. |
| RFX6 | -2.646949422 | -2.298023559 | -1.11566303 | 2.52E-19 | 5.95636E-19 | D.Q. | D.Q. | D.Q. |
| HSPA1A | 0.890722178 | 0.775074356 | -2.780692601 | 8.56E-61 | 6.06982E-60 | D.Q. | D.Q. | D.Q. |
| TMEM179 | 0.928963123 | 1.84540093 | -1.031097355 | 3.27E-17 | 7.085E-17 | D.Q. | D.Q. | D.Q. |
| PGBD5 | -0.110828622 | 0.6634803 | -1.524440899 | 5.05E-08 | 7.72353E-08 | D.Q. | D.Q. | D.Q. |
| ACSL3 | 0.001190277 | 0.980578875 | -1.299650064 | 7.53E-23 | 1.89465E-22 | D.Q. | D.Q. | D.Q. |
| REC8 | -0.746172319 | -1.753959417 | -1.435677882 | 4.04E-11 | 7.50286E-11 | D.Q. | D.Q. | D.Q. |
| LMO1 | 1.437365975 | 1.320263262 | -1.129046488 | 1.1E-41 | 5.72E-41 | D.Q. | D.Q. | D.Q. |
| FAM174B | 0.610566312 | 1.198162597 | -1.128520897 | 4.25E-36 | 1.84167E-35 | D.Q. | D.Q. | D.Q. |
| CRB3 | 1.488860713 | 2.19634992 | -1.66796438 | 2.52E-49 | 1.512E-48 | D.Q. | D.Q. | D.Q. |
| RGS11 | 1.721341105 | 1.654458601 | -2.954196845 | 5.24E-38 | 2.40424E-37 | D.Q. | D.Q. | D.Q. |
| WRAP73 | 0.398934572 | -0.685159461 | -1.237696076 | 1.24E-08 | 2.015E-08 | D.Q. | D.Q. | D.Q. |
| FRZB | -1.92322865 | -1.662622198 | -1.19486196 | 0.015072443 | 0.017038414 | D.Q. | D.Q. | D.Q. |
| DIRAS1 | 0.845862187 | 0.764335628 | -1.711547402 | 3.02E-12 | 5.74537E-12 | D.Q. | D.Q. | D.Q. |
| AARD | -1.209722256 | -1.193643721 | -1.479640348 | 0.0000789 | 0.000100889 | D.Q. | D.Q. | D.Q. |
| STK32A | 0.652621468 | 0.928806902 | -3.35870864 | 0.0000253 | 3.34475E-05 | D.Q. | D.Q. | D.Q. |
| SNCB | 0.070315831 | -0.7062394 | -1.627136024 | 2.88E-28 | 8.9856E-28 | D.Q. | D.Q. | D.Q. |
| PCDH10 | -1.324697374 | -1.146650109 | -1.23288173 | 0.0829545 | 0.087438527 | D.Q. | D.Q. | D.Q. |
| SHC3 | -0.756168302 | 0.687391774 | -1.070024019 | 0.149605061 | 0.151547984 | D.Q. | D.Q. | D.Q. |
| MAP7 | -0.819259154 | 1.1440893 | -1.198307089 | 1.04E-18 | 2.38588E-18 | D.Q. | D.Q. | D.Q. |
| MT3 | 0.611119786 | 1.183166232 | -2.713279068 | 0.01734379 | 0.019053741 | D.Q. | D.Q. | D.Q. |
| DPH1 | 1.089295226 | 1.00269326 | -1.693126543 | 0.007902084 | 0.009338827 | D.Q. | D.Q. | D.Q. |
| DISP2 | -0.145966122 | -1.067579622 | -1.310451006 | 0.004682441 | 0.005618929 | D.Q. | D.Q. | D.Q. |
| ESRP1 | 0.88383969 | 0.874622775 | -1.837212492 | 6.62E-32 | 2.34709E-31 | D.Q. | D.Q. | D.Q. |
| LIN7B | 1.065763058 | 1.987982055 | -1.854335479 | 2.28E-87 | 3.5568E-86 | D.Q. | D.Q. | D.Q. |
| TMEM179B | 1.392363081 | 0.852729062 | -1.279951704 | 1.89E-52 | 1.2285E-51 | D.Q. | D.Q. | D.Q. |
| SCO2 | 1.525666181 | 1.619435365 | -1.107320513 | 3.92E-10 | 7.1107E-10 | D.Q. | D.Q. | D.Q. |
| SLC7A4 | -0.205661147 | -1.015267 | -1.412493851 | 0.01334269 | 0.01530485 | D.Q. | D.Q. | D.Q. |
| KIRREL2 | -1.562025732 | -1.334690205 | -1.86528101 | 0.000000128 | 0.000000192 | D.Q. | D.Q. | D.Q. |
| SNX22 | 1.330621465 | 1.09543548 | -1.241135936 | 0.026422261 | 0.028624116 | D.Q. | D.Q. | D.Q. |
| ADAT3 | 0.689814745 | 1.435262289 | -1.774568136 | 0.099767508 | 0.103758208 | D.Q. | D.Q. | D.Q. |
| JPH3 | 0.822252042 | 2.086862665 | -1.076840018 | 0.000406734 | 0.000503575 | D.Q. | D.Q. | D.Q. |
| PRSS16 | 1.632900636 | 2.262997034 | -1.731366733 | 0.002811701 | 0.003426761 | D.Q. | D.Q. | D.Q. |
| SLC10A3 | 1.330203565 | 1.761527144 | -1.45191265 | 3.91E-13 | 7.6245E-13 | D.Q. | D.Q. | D.Q. |
| CHCHD10 | 1.815028389 | 2.020672255 | -1.123483675 | 9.27E-111 | 3.6153E-109 | D.Q. | D.Q. | D.Q. |
| UBL4A | 1.349697526 | 0.956438531 | -1.424353129 | 3.04E-16 | 6.24E-16 | D.Q. | D.Q. | D.Q. |
| CLSTN2 | -0.110398386 | -1.387419399 | -2.18558159 | 0.0000299 | 0.00003887 | D.Q. | D.Q. | D.Q. |
| VSTM2L | -1.990106796 | -1.414367985 | -1.025680011 | 0.008059312 | 0.009382483 | D.Q. | D.Q. | D.Q. |
| MAOB | -1.716323173 | -1.784122902 | -1.118275492 | 0.035163596 | 0.037572061 | D.Q. | D.Q. | D.Q. |
| GALNT18 | -1.227465074 | -0.724882101 | -2.339016463 | 3.01E-09 | 5.10391E-09 | D.Q. | D.Q. | D.Q. |
| SCRT1 | -0.084684974 | 0.77188793 | -2.827297554 | 1.54E-25 | 4.29E-25 | D.Q. | D.Q. | D.Q. |
| ATCAY | 0.31370842 | 1.431239112 | -2.058892807 | 0.015951283 | 0.017774287 | D.Q. | D.Q. | D.Q. |
| RNF213 | -0.111363257 | 0.964240974 | -1.695421212 | 6.37E-23 | 1.6562E-22 | D.Q. | D.Q. | D.Q. |
| BARX2 | -0.648042599 | -1.790371247 | -1.402599358 | 0.162572403 | 0.162572403 | D.Q. | D.Q. | D.Q. |
| SLC7A14 | -1.404923 | -0.937116452 | -1.463321066 | 0.000000424 | 6.01309E-07 | D.Q. | D.Q. | D.Q. |
| SNURF | -0.239693938 | 0.62462891 | -2.37316994 | 7.13E-17 | 1.50308E-16 | N.D. | N.D. | N.D. |
| SYNJ2BP | 0.171436074 | 1.00821525 | -1.057005749 | 1.89E-14 | 3.78E-14 | N.D. | N.D. | N.D. |
| CPNE1 | 0.316957978 | 1.103626392 | -1.21608947 | 3.54E-21 | 8.62875E-21 | N.D. | N.D. | N.D. |
| MRPS6 | 1.373071857 | 1.347072265 | -1.499210299 | 5.67E-33 | 2.2113E-32 | N.D. | N.D. | N.D. |
| HSPA5 | 1.643354748 | 1.947880412 | 0.753502516 | D.Q. | D.Q. | -0.629630073 | 4.86572E-06 | 3.34475E-05 |
