## Supplemental Table 4 for "Adaptation to chronic ER stress enforces pancreatic β-cell plasticity"

Supplementary table 4. Patients’ information in microarray analysis, related to Figure 6

| Sample ID | Clinical phenotype | Age (years) | Sex | Race | BMI | HLA Risk | Diabetes duration (years) | Source |
| --- | --- | --- | --- | --- | --- | --- | --- | --- |
| 6013 | Control | 65 | M | Caucasian | 24.2 | Neutral | N/A | nPOD |
| 6024 | Control | 21 | M | Caucasian | 27.8 | Predisposition | N/A | nPOD |
| 6048 | Control | 30 | M | Caucasian | 20.6 | Protective | N/A | nPOD |
| 6075 | Control | 16 | M | African American | 14.9 | Neutral | N/A | nPOD |
| 6012 | Control | 68 | F | Caucasian | 23.7 | Protective | N/A | nPOD |
| 6099 | Control | 14.2 | M | Caucasian | 30 | Predisposition | N/A | nPOD |
| 6140 | Control | 38 | M | Caucasian | 21.7 | Predisposition | N/A | nPOD |
| 6162 | Control | 22.7 | M | African American | 28.9 | Neutral | N/A | nPOD |
| 6168 | Control | 51 | M | Hispanic/Latino | 25.2 | Predisposition | N/A | nPOD |
| 6227 | Control | 17 | F | Caucasian | 26.4 | Predisposition | N/A | nPOD |
| 6129 | Control | 42.9 | F | Caucasian | 23.4 | Protective | N/A | nPOD |
| 6165 | Control | 45.8 | F | Caucasian | 25 | Protective | N/A | nPOD |
| 6102 | Control | 45.1 | F | Caucasian | 35.1 | Predisposition | N/A | nPOD |
| 6229 | Control | 31 | F | Caucasian | 26.9 | Neutral | N/A | nPOD |
| 6251 | Control | 33 | F | Caucasian | 29.5 | Neutral | N/A | nPOD |
| 6010 | Control | 47 | F | Caucasian | 19.7 | Protective | N/A | nPOD |
| 6179 | Control | 21.8 | F | Caucasian | 20.7 | Predisposition | N/A | nPOD |
| 6019 | Control | 42 | M | Caucasian | 31 | Predisposition | N/A | nPOD |
| 6080 | Antibody+ | 69.2 | F | Caucasian | 21.3 | Neutral | N/A | nPOD |
| 6123 | Antibody+ | 23.2 | F | Caucasian | 17.6 | Neutral | N/A | nPOD |
| 6158 | Antibody+ | 40.3 | M | Caucasian | 29.7 | Neutral | N/A | nPOD |
| 6167 | Antibody+ | 37 | M | Caucasian | 26.3 | Predisposition | N/A | nPOD |
| 6171 | Antibody+ | 4.3 | F | Caucasian | 14.8 | Predisposition | N/A | nPOD |
| 6044 | Antibody+ | 41.4 | M | Hispanic/Latino | 27.4 | Protective | N/A | nPOD |
| 6101 | Antibody+ | 64.8 | M | Caucasian | 34.3 | Protective | N/A | nPOD |
| 6154 | Antibody+ | 48.5 | F | Caucasian | 24.5 | Protective | N/A | nPOD |
| 6156 | Antibody+ | 40 | M | Caucasian | 19.8 | Neutral | N/A | nPOD |
| 6181 | Antibody+ | 31.9 | M | Caucasian | 21.9 | Predisposition | N/A | nPOD |
| 6197 | Antibody+ | 22 | M | African American | 28.2 | Neutral | N/A | nPOD |
| 6147 | Antibody+ | 23.8 | F | Caucasian | 32.9 | Neutral | N/A | nPOD |
| 6070 | T1D | 22.6 | F | Caucasian | 21.6 | Neutral | 7 | nPOD |
| 6084 | T1D | 14.2 | M | Caucasian | 26.3 | Predisposition | 4 | nPOD |
| 6088 | T1D | 31.2 | M | Caucasian | 27 | Predisposition | 5 | nPOD |
| 6180 | T1D | 27.1 | M | Caucasian | 25.9 | Predisposition | 11 | nPOD |
| 6224 | T1D | 21 | F | Caucasian | 22.8 | Predisposition | 1.5 | nPOD |
| 6243 | T1D | 13 | M | Caucasian | 21.3 | Predisposition | 5 | nPOD |
| 6228 | T1D | 13 | M | Caucasian | 17.4 | Predisposition | 0 | nPOD |
| 6209 | T1D | 5 | F | Caucasian | 11.95 | Predisposition | 0.25 | nPOD |
| 6038 | T1D | 37.2 | F | Caucasian | 30.9 | Predisposition | 20 | nPOD |
| 6046 | T1D | 18.8 | F | Caucasian | 25.2 | Predisposition | 8 | nPOD |
| 6069 | T1D | 22.9 | M | African American | 28.8 | Predisposition | 7 | nPOD |
| 6195 | T1D | 19.2 | M | Caucasian | 23.7 | Neutral | 5 | nPOD |
| 6052 | T1D | 12 | M | African American | 20.3 | Neutral | 1 | nPOD |
| 6268 | T1D | 12 | F | Caucasian | 26.6 | Predisposition | 3 | nPOD |
| 6265 | T1D | 11 | M | Caucasian | 12.9 | Predisposition | 8 | nPOD |
| 6196 | T1D | 26 | F | African American | 26.6 | Neutral | 15 | nPOD |
| 6211 | T1D | 24 | F | African American | 24.4 | Predisposition | 4 | nPOD |
| 6113 | T1D | 13.1 | F | Caucasian | 24.8 | Predisposition | 1.58 | nPOD |
| 6264 | T1D | 12 | F | Caucasian | 22 | Predisposition | 9 | nPOD |
| 6235 | T1D | 43.5 | M | Caucasian | 28.7 | Neutral | 21 | nPOD |
| 6188 | T2D | 36.1 | M | Unknown | Unknown | Unknown | Unknown | nPOD |
| 6114 | T2D | 42.8 | M | Unknown | Unknown | Unknown | Unknown | nPOD |
| 6249 | T2D | 45 | F | Unknown | Unknown | Unknown | Unknown | nPOD |
| 6275 | T2D | 48 | M | Unknown | Unknown | Unknown | Unknown | nPOD |
| 6273 | T2D | 45 | F | Unknown | Unknown | Unknown | Unknown | nPOD |
| 6191 | T2D | 62 | F | Unknown | Unknown | Unknown | Unknown | nPOD |
| 6059 | T2D | 18.8 | F | Unknown | Unknown | Unknown | Unknown | nPOD |
| 6110 | T2D | 20.7 | F | Unknown | Unknown | Unknown | Unknown | nPOD |
