## Supplemental Table 5 for "Adaptation to chronic ER stress enforces pancreatic β-cell plasticity"

Supplementary table 5. DNA oligo used in this study

| Name | oligo | Index | modifications | Note |
| --- | --- | --- | --- | --- |
| oCW-RTv2 | /5Phos/AGATCGGAAGAGCGTCGTGTAGGGAAAGAGTGT/iSp18/CAAGCAGAAGACGGCATACGAGATATTGATGGTGCCTACAG |  | 5'p, iSp18 | RT oligo |
| oCW-P7 | CAAGCAGAAGACGGCATACGAGAT |  |  | PCR |
| oCW1AD005v2 | AATGATACGGCGACCACCGAGATCTACACGATCGGAAGAGCACACGTCTGAACTCCAGTCACACAGTGACACTCTTTCCCTACACGACGC | ACAGTG |  | PCR |
| oCW1AD006v2 | AATGATACGGCGACCACCGAGATCTACACGATCGGAAGAGCACACGTCTGAACTCCAGTCACGCCAATACACTCTTTCCCTACACGACGC | GCCAAT |  | PCR |
| oCW1AD012v2 | AATGATACGGCGACCACCGAGATCTACACGATCGGAAGAGCACACGTCTGAACTCCAGTCACCTTGTAACACTCTTTCCCTACACGACGC | CTTGTA |  | PCR |
| oCW1AD019v2 | AATGATACGGCGACCACCGAGATCTACACGATCGGAAGAGCACACGTCTGAACTCCAGTCACGTGAAACCACTCTTTCCCTACACGACGC | GTGAAA |  | PCR |
| oCW2AD002v2 | AATGATACGGCGACCACCGAGATCTACACGATCGGAAGAGCACACGTCTGAACTCCAGTCACCGATGTACACTCTTTCCCTACACGACGC | CGATGT |  | PCR |
| oCW2AD004v2 | AATGATACGGCGACCACCGAGATCTACACGATCGGAAGAGCACACGTCTGAACTCCAGTCACTGACCAACACTCTTTCCCTACACGACGC | TGACCA |  | PCR |
| oCW2AD007v2 | AATGATACGGCGACCACCGAGATCTACACGATCGGAAGAGCACACGTCTGAACTCCAGTCACCAGATCACACTCTTTCCCTACACGACGC | CAGATC |  | PCR |
| oCW2AD016v2 | AATGATACGGCGACCACCGAGATCTACACGATCGGAAGAGCACACGTCTGAACTCCAGTCACCCGTCCCCACTCTTTCCCTACACGACGC | CCGTCC |  | PCR |
| oCW3AD001v2 | AATGATACGGCGACCACCGAGATCTACACGATCGGAAGAGCACACGTCTGAACTCCAGTCACATCACGACACTCTTTCCCTACACGACGC | ATCACG |  | PCR |
| oCW3AD008v2 | AATGATACGGCGACCACCGAGATCTACACGATCGGAAGAGCACACGTCTGAACTCCAGTCACACTTGAACACTCTTTCCCTACACGACGC | ACTTGA |  | PCR |
| oCW3AD010v2 | AATGATACGGCGACCACCGAGATCTACACGATCGGAAGAGCACACGTCTGAACTCCAGTCACTAGCTTACACTCTTTCCCTACACGACGC | TAGCTT |  | PCR |
| oCW3AD011v2 | AATGATACGGCGACCACCGAGATCTACACGATCGGAAGAGCACACGTCTGAACTCCAGTCACGGCTACACACTCTTTCCCTACACGACGC | GGCTAC |  | PCR |
| oCW4AD003v2 | AATGATACGGCGACCACCGAGATCTACACGATCGGAAGAGCACACGTCTGAACTCCAGTCACTTAGGCACACTCTTTCCCTACACGACGC | TTAGGC |  | PCR |
| oCW4AD009v2 | AATGATACGGCGACCACCGAGATCTACACGATCGGAAGAGCACACGTCTGAACTCCAGTCACGATCAGACACTCTTTCCCTACACGACGC | GATCAG |  | PCR |
| oCW4AD022v2 | AATGATACGGCGACCACCGAGATCTACACGATCGGAAGAGCACACGTCTGAACTCCAGTCACCGTACGTCACTCTTTCCCTACACGACGC | CGTACG |  | PCR |
| oCW4AD027v2 | AATGATACGGCGACCACCGAGATCTACACGATCGGAAGAGCACACGTCTGAACTCCAGTCACATTCCTTCACTCTTTCCCTACACGACGC | ATTCCT |  | PCR |
| mATF3_R | TTGACGGTAACTGACTCCAGC |  |  | RT-qPCR |
| mATF3_F | GAGGATTTTGCTAACCTGACACC |  |  | RT-qPCR |
| mATF4-F | GTTTGACTTCGATGCTCTGTTTC |  |  | RT-qPCR |
| mATF4-R | GGGCTCCTTATTAGTCTCTTGG |  |  | RT-qPCR |
| mERO1B-F | TGGACACTGGGCAAAGAGAC |  |  | RT-qPCR |
| mERO1B-R | ACAAAATGTGAAACAAACATGGCA |  |  | RT-qPCR |
| mGLUT2-F | ACACCGGAATGTTCTTAGCC |  |  | RT-qPCR |
| mGLUT2-R | GTGAGAGAAGCCGAGGAAAG |  |  | RT-qPCR |
| mHERPUD1-F | TGCTGTTGGATCACCAGTGT |  |  | RT-qPCR |
| mHERPUD1-R | CTGTGGATTCAGCACCCTTT |  |  | RT-qPCR |
| mINS1-F | CCTGTTGGTGCACTTCCTAC |  |  | RT-qPCR |
| mINS1-R | GTGTGTAGAAGAAGCCACGCTC |  |  | RT-qPCR |
| mINS2-F | GGAGCGTGGCTTCTTCTACA |  |  | RT-qPCR |
| mINS2-R | GGTCTGAAGGTCACCTGCTC |  |  | RT-qPCR |
| mMAFA-F | GAGGTCATCCGACTGAAACAGAAGC |  |  | RT-qPCR |
| mMAFA-R | TGGAGCTGGCACTTCTCGCT |  |  | RT-qPCR |
| mP4HB-F | GCTGTGCGGCTTATTACCCT |  |  | RT-qPCR |
| mP4HB-R | CTCATCAGGTGGGGCTTGAT |  |  | RT-qPCR |
| mPCSK2-F | AGAGGAATCCTGAGGCTGGT |  |  | RT-qPCR |
| mPCSK2-R | ATCCAGGTTAGCCTCCAAAGC |  |  | RT-qPCR |
